# Evolutionary dynamics of Respiratory Syncytial Virus in pre-pandemic, pandemic, and post-pandemic periods in Houston, Texas, USA

**DOI:** 10.1101/2025.08.06.668939

**Authors:** Vasanthi Avadhanula, Daniel P Agustinho, Leila C. Sahni, Anil Surathu, Shelby. R. Simar, David Henke, Harshavardhan Doddapaneni, Donna M Muzny, Ginger A Metcalf, Sara Javornik Cregeen, Natalie J. Thornburg, Heidi L. Moline, Ayzsa Tannis, Richard A Gibbs, Joseph F Petrosino, Julie A Boom, Fritz J Sedlazeck, Pedro A Piedra

## Abstract

**Background:** Respiratory syncytial virus (RSV) is a leading cause of severe lower respiratory tract infections in infants and poses significant risks to immuno-compromised individuals and the elderly. The COVID-19 pandemic disrupted typical RSV seasonality, leading to unusual patterns of viral circulation and resurgence. However, the impact of these disruptions on RSV evolutionary dynamics remains incompletely understood. This study aimed to investigate the evolutionary dynamics of RSV in Houston, Texas, across pre-pandemic (November 2015 – March 2020), pandemic (April 2021 – June 2023), and post-pandemic periods (July 2023 – December 2024), focusing on genetic diversity, lineage dynamics, and selective pressures.

**Methods:** In one of the largest comprehensive RSV genomic evolutionary analyses, we analyzed 1,344 RSV-positive respiratory samples collected from children seeking outpatient care or hospitalized with acute respiratory infections between November 2015 to February 2024, which is one of the largest comprehensive RSV genomic evolutionary analyses. We successfully sequenced the whole genomes for 87.5% of the isolates (606 RSV/A and 570 RSV/B). To assess genetic diversity, lineage dynamics, and selective pressures, we conducted phylogenetic analysis, variant calling, and dN/dS ratio calculations. We clustered variant prevalence patterns using k-means and used Shannon entropy to quantify genetic variability. We performed statistical regression analyses to evaluate the accumulation of variants and their impact on gene expression.

**Results:** RSV circulation in Houston was markedly disrupted during the pandemic, with an absence of cases in 2020-2021 followed by off-season surges from 2021 to 2023. Phylogenetic analysis revealed distinct lineage dynamics, with RSV/A showing remarkable persistence of specific lineages such as A.D.1 and emergence of new lineages including A.D.3. In contrast, RSV/B underwent a dramatic restructuring, dominated by the B.D.E.1 lineage in the pandemic and post-pandemic period. Bayesian molecular clock analysis estimated nucleotide substitution rates of 9.37 × 10^−4^ (95% HPD: 7.99 × 10^−4^ − 1.07 × 10^−3^) substitutions/site/year for RSV/A and 9.79 × 10^−4^ (95% HPD: 8.96 × 10^−4^ – 1.06 × 10^−3^) for RSV/B. The time to the most recent common ancestor (tMRCA) of the dominant RSV/A clade, A.D.1, is dated to approximately 2010 (95% HPD: 2008–2011), while for RSV/B the dominant clade- B.D.E.1 traces back to 2002 (95% HPD: 1997–2007). Despite reduced lineage diversity, RSV/B accumulated non-synonymous variants at nearly twice the rate of RSV/A, notably in the M2-2 gene, reflecting high within-lineage variability. The significant increase in non-synonymous mutations in M2-2 in RSV/B during the pandemic and post-pandemic time phases strongly correlated with increased transcriptional activity. Additionally, antigenic site mutations in the F protein, particularly in RSV/B, were also observed, with implications for immune evasion and effectiveness of the current monoclonal antibody immunoprophylaxis intervention.

**Conclusions:** The COVID-19 pandemic significantly impacted RSV evolution, leading to reduced genetic diversity during the pandemic and the emergence of novel lineages post-pandemic. RSV/B exhibited more dynamic evolutionary changes, particularly in the M2-2 gene, suggesting potential adaptive advantages. Ongoing genomic surveillance coupled with functional studies is crucial for monitoring variants and assessing the clinical impact of these mutations on transmission, disease severity, fitness, and long-term effectiveness of RSV prevention and treatment strategies.

## Background

Respiratory syncytial virus (RSV) is the leading cause of severe lower respiratory tract infections in infants worldwide, with nearly all children being infected by the age of two years (Glezen et al., 1986; Y. Li et al., 2021). RSV infection can recur throughout a person’s life, posing a risk of serious infections, particularly in the immunocompromised, and the elderly (Falsey et al., 2005). Recently, three RSV vaccines have been approved by the U.S. Food and Drug Administration and recommended by the Advisory Committee on Immunization Practices (ACIP) of the Centers for Disease Control and Prevention (CDC) for use in adults aged 75 and older and for those aged 18 to 74 who are at an increased risk of severe RSV (Alberto et al., 2023; E et al., 2023; Eleanor et al., 2023). In addition, three new immunization options are recommended to protect infants from severe RSV: a maternal RSV vaccine given during pregnancy at 32 to 36 weeks of gestation to safeguard the newborn up to 6 months of age and two different long acting RSV monoclonal antibodies (nirsevimab and clesrovimab) both can be administered to all infants <8 months of age born during or entering their first RSV season who were not protected by maternal RSV vaccination and to high-risk infants before their second RSV season (Beate et al., 2023; L et al., 2022).

RSV has a negative-sense, single-stranded, non-segmented genome with ∼15,200 nucleotides comprising 10 genes (3′ NS1-NS2-N-P-M-SH-G-F-M2-L 5′) that encode 11 proteins: non-structural proteins 1 and 2 (NS1 and NS2), the nucleocapsid protein (N), phosphoprotein (P), matrix protein (M), small hydrophobic protein (SH), attachment glycoprotein (G), fusion glycoprotein (F), the M2 gene encoding two proteins; transcription processivity factor M2-1 and the regulatory protein M2-2; and the large RNA-dependent RNA polymerase (L)(Collins & Melero, 2011). RSV comprises two major subgroups, RSV/A and RSV/B, which are distinguished by genetic and antigenic variations in their G glycoprotein (Anderson et al., 1985; Peret et al., 1998; Venter et al., 2001). Although RSV’s molecular epidemiology and evolutionary dynamics have been studied extensively, most analyses have primarily focused on the G and F genes (Hause et al., 2017; Trento et al., 2006; Venter et al., 2001). Below the subgroup level, RSV/A and RSV/B are further classified into lineages using a recently standardized hierarchical nomenclature system based on whole-genome phylogenetic analysis, analogous to the Pango nomenclature used for SARS-CoV-2(Goya et al., 2024). Limited information is available regarding the diversity and evolution rates of the overall RSV genome. A few studies have indicated that the surface glycoproteins SH and G, along with the accessory transcription regulation protein (M2-2), exhibit the highest levels of sequence variability among RSV/A and RSV/B whole genomes (Lydia et al., 2013; Schobel et al., 2016). Analyzing fluctuations in RSV’s whole genome over time can elucidate the roles of specific genes in RSV survival. Large longitudinal studies are therefore needed to uncover how genomic changes influence RSV epidemics and evolution, especially in the context of the newly approved RSV vaccines and monoclonal antibody interventions.

In the U.S., RSV outbreaks typically followed a seasonal pattern before the COVID-19 pandemic. In Houston, TX, the RSV season typically begins in October, peaks in December or January, and ends in April. However, the COVID-19 pandemic disrupted this pattern during 2020-2022, leading to an absence of RSV circulating in 2020-2021, followed by two years of unusual and prolonged RSV surges. There was an off-season resurgence in 2021-2022 and 2022-2023, with peaks occurring in July and November, respectively (Hamid et al., 2023). During the 2023-2024 season, RSV circulation had returned to the typical pre-pandemic pattern, starting in the fall and peaking in the winter (NVSN, 2024).

The atypical and prolonged RSV outbreaks during the 2021-2023 period resulted in a higher proportion of RSV hospitalizations and intensive care unit (ICU) admissions worldwide (Hatter et al., 2021; McMorrow et al., 2024; Ujiie et al., 2021). Genomic studies from various regions have provided insight into the reasons for this upsurge in RSV. Early data from Australia indicated a significant reduction in RSV genetic diversity following SARS-CoV-2 circulation, with two genetically distinct RSV/A lineages circulating cryptically and later causing extensive outbreaks and hospitalizations in early 2021 (Eden et al., 2022). A study from South Africa noted that the same pre-existing strains circulating globally were present during the 2021-2022 resurgence, and heightened infections in children were attributed to "immune debt" (also referred to as "immunity gap"), characterized by the lack of natural infections during the first year of the pandemic thereby rendering infants and young children susceptible at an older age (Sondlane et al., 2024). Conversely, studies from Italy, Japan, and Argentina indicated that new introductions of various lineages or distinct RSV/B lineages were responsible for these regional surges (Dolores et al., 2022; Korsun et al., 2024; Pierangeli et al., 2024; Simões et al., 2021). Despite these studies, the evolutionary dynamics of RSV during the pre-pandemic, pandemic, and post-pandemic periods have not been fully explored. Investigating these dynamics would offer valuable insights into how isolates from the same geographic location differ across these distinct time frames.

The rapid evolution of RSV is driven by selective immune pressures from the host, resulting in antigenic variations over time that can significantly impact the effectiveness of antiviral treatments, monoclonal antibody intervention, and vaccines. There have been reports of community-acquired mutations or prophylactic escape mutants in the F protein, which is a primary target of monoclonal antibodies and vaccines. For example, phase III clinical trials of the monoclonal antibody suptavumab were unsuccessful due to the community-wide emergence of resistant mutations in the F protein of RSV/B strains, which reduced the antibody’s binding affinity(Simões et al., 2021). Mutations associated with palivizumab, the first approved monoclonal antibody for immunoprophylaxis, have been reported only sporadically (Adams et al., 2010; Zhu et al., 2011), likely due to its limited use in a small subset of high-risk infants. Currently, there is no evidence that the 2024 recommendation of the monoclonal antibody nirsevimab for all infants not protected by maternal RSV vaccination will create additional selective pressure on the virus. However, monitoring RSV’s molecular evolution and the impact of vaccines and monoclonal antibodies is essential, especially during this period of viral evolution and emerging interventions.

In this study, we investigated the genetic diversity and evolutionary dynamics of RSV in children residing in the greater Houston metropolitan area over a nine-year period from 2015 to 2024, which encompassed the pre-pandemic, pandemic, and post-pandemic periods. The samples analyzed were collected from a single geographic location: Houston, Texas, as part of the New Vaccine Surveillance Network (NVSN)(CDC, 2025).

NVSN is a prospective, active, population-based surveillance platform enrolling children at seven pediatric medical centers who present with acute respiratory illnesses (ARI). In this comprehensive analysis of 1,344 RSV isolates from Houston from 2015 to 2024, we observed distinct evolutionary patterns between RSV subgroups RSV/A and RSV/B. RSV/B exhibited reduced lineage diversity during the pandemic and post-pandemic periods, even though it accumulated a higher number of non-synonymous variants than RSV/A, especially in SH, F & M2-2 genes. The increase in RSV/B M2-2 variants correlated significantly with heightened transcript levels in all 10 RSV genes, and possibly providing fitness advantages. For both subgroups, the F protein continued to accumulate non-synonymous variants, warranting continued surveillance of RSV genomes, given newly approved vaccines, and long-acting monoclonal antibodies target the F protein..

## Methods

### Study Design

The study was conducted within the NVSN, a Centers for Disease Control and Prevention (CDC)-led network of seven locations in the U.S. that performs population-based surveillance for ARI in children. For this analysis, we used data from only the Houston site. Children 0 to <18 years presenting with ARI were enrolled year-round in the outpatient or urgent care clinics, emergency department (ED), or hospital between December 5, 2015, and February 15, 2024. Mid-turbinate nasal and/or oropharyngeal flocked swabs were obtained; if both were collected, they were combined and placed together in a universal transport medium. Tracheal aspirates were obtained in the 2015-16 season only and were an alternative specimen for patients who were intubated. If no sample was obtained during the visit, a clinical leftover salvage sample was used, which may have included various nasal swab types or nasal washes. The study received Institutional Review Board approval from Baylor College of Medicine and the CDC.

To categorize samples temporally, we divided the collection dates into three distinct time phases: "Pre-pandemic" (November 4, 2015, to March 20, 2020), "Pandemic" (April 7, 2021, to June 30, 2023), and "Post-pandemic" (July 1, 2023, to December 12, 2024). These time phases were defined to capture potential variations in RSV epidemiology before, during, and after the COVID-19 pandemic period. Of the 94 samples collected between July 1, 2023, to December 12, 2024, during the “post-pandemic period,” 4 infants had an RSV breakthrough infection after receiving nirsevimab monoclonal antibody.

### Real-time PCR testing

The detection of RSV was performed using a real-time reverse-transcription polymerase chain reaction (RT-PCR). RSV/A and RSV/B were detected by one-step RT-PCR, as described previously (Avadhanula et al., 2015; Perez et al., 2022).

### cDNA preparation for whole genome sequencing of RSV

cDNA preparation was done as previously described (Avadhanula et al., 2024; Bhamidipati et al., 2025). RNA was analyzed using the RNA 6000 Nano assay (Agilent) or the RNA 6000 Pico assay for determination of an RNA Integrity Number (RIN) and DV200 metrics. Strand specific cDNA was generated utilizing NEBNext® RNA First Strand Synthesis Module (E7525L; New England Biolabs Inc.) and NEBNext® Ultra™ II Directional RNA Second Strand Synthesis Module (E7550L; New England Biolabs Inc.), which produces strand-specific libraries by incorporating dUTP during second-strand synthesis. This directionality preserves the strand origin of RNA molecules and enables distinction between genomic RNA (negative sense) and mRNA transcripts (positive sense) in downstream analyses. Total RNA in a 15 μL mixture containing random primers and 2X 1st strand cDNA synthesis buffer was incubated at 94 °C for 10 min to fragment the RNA to 200-600bp. RNA was converted to cDNA by adding a 5 µL enzyme mix containing 500 ng Actinomycin D (A7592, Thermo Fisher Scientific), 0.5 μLRNase inhibitor, and 1 μL of Protoscript II reverse transcriptase, then incubated at 25 °C for 10 minutes, 42 °C for 50 minutes, 70 °C 15 minutes, before being cooled to 4 °C on a thermocycler. Second-strand cDNA was synthesized by adding 60 μL of the mix containing 48 μL H2O, 8 μL of 10X reaction buffer, and 4 μL of second-strand synthesis enzyme, and incubating at 16 °C for 1 hour on a thermocycler. The double-stranded (ds) cDNA was purified with 1.8X volume of AMPure XP beads (A63882, Beckman) and eluted into 42 μL 10 mM Tris buffer (Cat#A33566, Thermo Fisher Scientific). Because these libraries were prepared primarily for sequence capture, rRNA depletion, or Poly A+ RNA isolation steps were not performed.

### Library preparation for whole genome sequencing of RSV

Pooled cDNA libraries were hybridized with biotin-labeled probes from the RSV Panel (Twist Biosciences, Inc) at 70 °C for 16 hours as previously described (Avadhanula et al., 2024; Bhamidipati et al., 2025). The RSV probe set size was 23.77 Mb and was designed based on 1,570 publicly available genomic sequences of RSV isolates. In this probe set, there are 87,025 unique probes of 80 bp length, which cover 99.79 percent of the targeted isolates. Captured virus targets were incubated with streptavidin beads for 30 min at room temperature. Streptavidin beads bound with virus targets were washed and amplified with the KAPA HiFi HotStart enzyme. The amount of each cDNA library pooled for hybridization and post-capture amplification PCR cycles (12 ∼ 20) was determined empirically according to the virus Ct values. In general, between 1.8–4.0 μg pre-capture library was used for hybridization with the probes, and the post-capture libraries were sequenced on Illumina NovaSeq S4 flow cell, to generate 2 × 150 bp paired-end reads.

### Reference genome

To facilitate comparisons between RSV/A and RSV/B subgroups, we generated consensus reference genomes, NewRSVA2014 and NewRSVB2014, respectively, from the genotypes in circulation, RSV/A/Ontario (RSV/A/ON) and RSV/B/Buenos Aires (RSV/B/BA). These were constructed using 53 RSV/A/ON and 121 RSV/B/BA strain sequences collected in 2014 from multiple regions of the world, the year preceding our sample collection period (details in Supplementary Table 1). To create the consensus, FASTA sequences were processed by a customized RSV module within Iterative Refinement Meta-Assembler v. 0.9.3 (IRMA-RSV)(Shepard et al., 2016). IRMA-RSV was created by constructing the reference assembly templates for different subgroups and genotypes of RSV from publicly available nucleotide sequences. Briefly, both RSV/A/ON and RSV/B/BA whole-genome consensus sequences were retrieved from NCBI’s Nucleotide database, aligned with MUSCLE v. 3.8.3130303030 and built with the maximum likelihood phylogeny in RAxML v8.2 (Stamatakis, 2014) with an autoMRE option, for an efficient and automatic bootstrapping convergence criterion. The reference genomes are available as FASTA files. The list of RSV isolates and their accession numbers are listed in Supplementary Table 1.

We annotated the reference genomes using Vigor4 (Wang et al., 2012), a specialized viral genome annotation tool. The annotation process utilized a pre-defined RSV-specific protein sequence database downloaded from the Vigor4 reference GitHub repository (https://github.com/JCVenterInstitute/VIGOR_DB/blob/master/Reference_DBs/rsv_db). We generated the genome annotation by running Vigor4 on the consensus reference genomes and applying the RSV-specific database to identify and characterize viral genes. The resulting genome annotation file in GFF3 format was subsequently converted to GTF format using gffread from the Cufflinks package (Pertea & Pertea, 2020), enabling compatibility with downstream genomic analyses.

### Genome alignment and variant calling

After sequencing, adaptors were identified and removed, and reads were trimmed for quality using Trimmomatic version 0.39–2 (Bolger et al., 2014) with the following parameters: ‘ILLUMINACLIP:TruSeq3-PE.fa:2:30:10:2:keepBothReads LEADING:3 TRAILING:3 MINLEN:36’. Trimmed reads were taxonomically classified using Kraken2 version 2.1.2 (Wood et al., 2019) with default parameters. Reads were filtered using the Kraken2 script extract_kraken_reads.py used with the parameters ‘-t 11244 -include-children’, which ensures that only reads classified as belonging to the Pneumoviridae family are kept. To independently assess the genotype of each sample, the RSV/A/ON and RSV/B/BA reference genomes were concatenated, and the resulting FASTA file was used as a reference genome for BWA-MEM version 0.7.17-r1188 (H. Li & Durbin, 2010). The resulting SAM files were converted to BAM, sorted, and indexed using samtools version 1.6(H. Li et al., 2009). To detect variants on the BAM files, samtools mpileup (1.6) was used with the ‘-aa -A -d 0 -B -Q 0’ parameters, followed by iVar version 1.3.1(Grubaugh et al., 2019) with standard parameters, using the same reference genome as the one used for alignment. The resulting TSV files generated by iVar were converted to VCF using the script ivar_variants_to_vcf.py from iVar. Variant annotation was performed using SnpEff version 4.3 u (Cingolani et al., 2012). In this study, we defined a "variant" as a nucleotide difference relative to the consensus reference genome, identified at a minimum depth of 30 reads and an alternative allele frequency of ≥20%. These represent population-level consensus changes rather than intra-host single nucleotide polymorphisms (iSNPs). Variants were filtered using bcftools view(Danecek et al., 2021). Plots and subsequent analysis were generated using the R package *ggplot*2.

### Sample Quality Control and Genotype Assignment

To ensure high-quality genomic data for downstream analyses, samples underwent stringent quality control filtering based on breadth of coverage. Coverage depth was calculated for both RSV/A and RSV/B reference genomes, and the proportion of each genome covered at ≥20x depth was determined. Samples were included in subsequent analyses only if they met a minimum breadth of coverage threshold of 90% (i.e., ≥90% of the genome had coverage ≥20x).

In silico genotyping was performed based on these coverage metrics and can be viewed in Supplementary Table 2. Samples with ≥90% breadth of ≥20x coverage for RSV/A were classified as RSV/A infections, while those meeting the same criterion for RSV/B were classified as RSV/B infections. Samples meeting the coverage threshold for both genotypes were classified as co-infections, while those failing to meet the 90% threshold for either genotype were classified as low-coverage infections. Post hoc inspection of the BEAST2 time-calibrated phylogeny for RSV/A identified two additional samples with detectable RSV/B reads that had passed the initial 90% breadth-of-coverage filter. These co-infected samples produced chimeric consensus sequences that were phylogenetically divergent. These two samples were excluded from the molecular clock analysis and time-calibrated tree figures. Both co-infections and low-coverage samples were excluded from all downstream analyses to ensure data quality and avoid ambiguous genotype assignments.

### Genome assembly and lineage classification

Following sequencing, raw data files in binary base call (BCL) format were converted into FASTQs and demultiplexed based on the dual-index barcodes using the Illumina ‘bcl2fastq’ software. Demultiplexed raw FASTQ sequences were processed using BBDuk to quality trim, remove Illumina adapters, and filter PhiX reads. Trimmed FASTQs were mapped to a combined PhiX and human reference genome database (GRCh38) using BBMap to determine and remove human/PhiX reads. Trimmed and host-filtered reads were processed through VirMAP (Ajami et al., 2018) to assemble full-length RSV genomes. The VirMAP summary statistics include information on reconstructed genome length, the number of reads mapped to the reconstruction, and the average coverage across the genome. Final reconstructions were manually inspected using Geneious Prime® 2022.1.1 and aligned against the relevant RSV reference genomes to determine the quality of assemblies.

For lineage classification, we uploaded a multi-fasta file containing all RSV/A or RSV/B assembled genomes to UShER (https://genome.ucsc.edu/cgi-bin/hgPhyloPlace) (Turakhia et al., 2021) to assign them to a lineage as described by Goya et al. (Goya et al., 2024). The RSV lineage (listed as PANGO lineage in the software’s output) and Nextstrain clade information generated were added to our metadata and added to the phylogenetic analysis performed, as described below.

### Phylogenetic analysis

To reconstruct the phylogenetic relationships among RSV isolates, we first generated alignments directly from raw sequencing reads using Omni2Tree (Majidian et al., 2026). The resulting concatenated amino acid alignment was then used for phylogenetic inference using IQ-TREE (version 2.2.0.3). To assess the statistical support for the tree topology, we performed ultrafast bootstrap approximation with 1,000 replicates (-bb 1000). The final tree with bootstrap support values was visualized using the *ggtree* R package (G. Yu et al., 2017).

Phylogenetic relationships among RSV isolates were interpreted using the standardized RSV lineage nomenclature(Goya et al., 2024) assigned via UShER placement on the global reference phylogeny. Clades within the local phylogeny were described by their lineage composition and temporal distribution across the three time- phases.

### Molecular Clock Analysis

To estimate evolutionary rates and divergence times, we performed Bayesian phylogenetic inference using BEAST v2.6.3(Bouckaert et al., 2019) on whole-genome nucleotide alignments, analyzed separately for RSV/A (n=659) and RSV/B (n=693). We employed the HKY+Γ₄ substitution model to account for transition/transversion bias and among-site rate heterogeneity. Given the potential for rate variation across viral lineages driven by differential immune selection pressures, we implemented an uncorrelated relaxed molecular clock with rates drawn from a lognormal distribution(Drummond et al., 2006). We specified a coalescent constant population size tree priori, appropriate for endemic viruses with relatively stable transmission dynamics. Tip dates were specified using decimal year sampling dates encoded in sequence names. To ensure adequate sampling of the posterior distribution, we ran multiple independent Markov chain Monte Carlo (MCMC) chains per subtype with different random seeds, each for up to 200–500 million iterations, sampling every 1,000 steps. Chains were combined using LogCombiner v2.6.3 with 10% burn-in, and convergence was assessed using Tracer v1.7.2(Rambaut et al., 2018), ensuring effective sample sizes (ESS) >200 for all parameters. The combined posterior samples were resampled to approximately 14,000 trees per subtype for computational tractability. Maximum clade credibility (MCC) trees were summarized using TreeAnnotator v2.6.3 with common ancestor node heights and visualized using the ggtree R package(G. Yu et al., 2017). BEAST2 molecular clock parameter estimates can be seen in **Supplementary Table 4**.

### Statistical analysis

All scripts and analysis details used in this study are available on GitHub at https://github.com/DanielPAagustinho/RSV_pandemics. To account for potential subgroup-specific evolutionary patterns, all analyses were conducted separately for RSV/A and RSV/B. We included variants from all 11 RSV genes (NS1, NS2, N, P, M, SH, G, F, M2-1, M2-2, and L) and considered all effect types in the dataset. Unless otherwise stated, all analyses were conducted using R (version 4.2.1).

### Statistical analysis of variant accumulation

To evaluate the accumulation of RSV genetic variants over time, a linear regression model was employed, accounting for both RSV/A and RSV/B strains. The model was specified as:

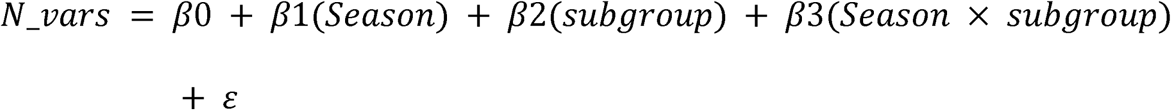

where β0 represents the baseline number of variants, β1 quantifies the rate of variant accumulation per season for RSV/A, β2 estimates the difference in baseline variants between RSV subgroups, and β3 tests whether RSV/B accumulates variants at a different rate than RSV/A. Seasons were coded numerically from 1 (2015-2016) to 7 (2023-2024). A similar model was applied to all variants, all non-synonymous variants, and non-synonymous variants per gene. This interaction model is a standard approach for comparing slopes between groups in linear regression(Aiken, 1991).

### Variant prevalence behavior clustering

To analyze the evolutionary dynamics of RSV variants, we applied k-means clustering to group variants by their prevalence behavior across time phases. We constructed a matrix where each row represented a unique variant and the three columns represented the scaled prevalence (population allele frequency) of that variant in the pre-pandemic, pandemic, and post-pandemic periods, respectively. Prevalence values were z-score scaled to ensure equal weighting of each time phase. The k-means algorithm (stats package, base R) was then applied to this three-dimensional feature space. We selected eight clusters (k = 8) based on elbow-method analysis and visual inspection of cluster stability, which provided biologically interpretable groupings of variant behavior.

### dN/dS calculation

We calculated the ratio of nonsynonymous to synonymous substitution rates (dN/dS) for each combination of season, viral subgroup (RSV/A and RSV/B), and gene to investigate selective pressures acting on viral genes. After filtering out upstream and downstream gene variants, we computed the rates as follows:

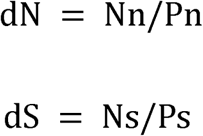

where Nn is the total count of nonsynonymous variants, Pn is the number of unique positions with nonsynonymous changes, Ns is the total count of synonymous variants, and Ps is the number of unique positions with synonymous changes. The final dN/dS ratio was then calculated as:

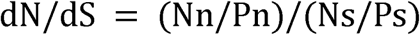

In cases where no synonymous variants were observed (dS = 0), dN was set to zero to avoid undefined ratios.

The individual ratio of non-synonymous to synonymous variants for all genes was calculated by dividing the number of non-synonymous variants by the number of synonymous variants for each isolate.

### Calculation of Negative Strand Ratios

We previously demonstrated the use of probe-based capture enrichment sequencing to measure both genomes and antigenomes of RSV utilizing strand-specific libraries (Bhamidipati et al., 2025). (Liao et al., 2014). Because our library preparation was strand-specific (see cDNA preparation above), we used the strand orientation of mapped reads as a proxy for the relative abundance of genomic RNA versus mRNA transcripts, as previously described(Bhamidipati et al., 2025). RSV is a negative-sense RNA virus; therefore, in a strand-specific library, reads derived from mRNA transcripts (positive sense) map to the antisense strand of the reference genome, while reads from genomic RNA map to the sense strand. We quantified strand-specific counts using featureCounts(Liao et al., 2014) with the -s 1 parameter for sense-strand counts and -s 2 for antisense-strand counts. For each gene, we calculated the transcript ratio as the proportion of antisense strand reads (representing mRNA) relative to total reads (sense + antisense). This approach allowed us to assess transcriptional activity patterns and their relationship to specific genetic variants across RSV isolates.

### Shannon entropy calculation

To quantify genetic variability across viral genes, we calculated Shannon entropy for each gene separately in RSV subgroups A and B (Hause et al., 2017). We used the multiple sequence alignment generated by Omni2Tree, containing consensus sequences for each isolate.

The Shannon entropy calculation involved transforming sequence alignments through several key steps. First, we removed reference strains from the alignment. Then, we converted the sequences into a character matrix and calculated the frequency of each amino acid at each position. We computed Shannon entropy using the formula:

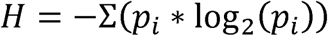

where p_i_ represents the frequency of each amino acid at a given position.

## Results

### RSV epidemiology and sequencing of the cohort

Over nine years (2015-2024), we collected a total of 11,396 respiratory samples from children presenting with ARI. These samples underwent routine RSV testing by RT-PCR. Of these samples, 1,020 (8.9%) tested positive for RSV/A, 1,109 (9.7%) for RSV/B, and 13 for both RSV subgroups. The monthly distribution of positive cases is presented in **Suppl. Fig. 1A**. Seasonal circulation patterns of RSV in Houston mirrored the patterns observed in the rest of the US (Hamid et al., 2023). In the pre-pandemic time phase, RSV circulated during the fall and winter months. This was followed by a near-complete absence of RSV cases from March 2020 to May 2021 during the first year of the COVID-19 pandemic, with only sporadic detections during this period. When nonpharmaceutical interventions (NPIs) were lifted, RSV resurged out of season in May 2021, followed by another summer peak in 2022. RSV circulation returned to the pre-pandemic pattern of fall/winter months in the 2023-24. The RSV seasons were either dominated by a single RSV subgroup or showed a relatively equal representation of both subgroups.

To explore the genetic makeup of circulating RSV strains, we selected a subset of 1,344 (62.7%) samples with high viral loads (CT ≤30) for whole-genome sequencing. This comprised 654 (64.1%) RSV/A positive samples, 683 (61.6%) RSV/B positive samples, and 7 (53.8%) RSV/A and RSV/B co-infection samples (**Suppl. Fig. 1B**). Upon sequencing and filtering for coverage and quality (see Methods), we retained 606 RSV/A (92.6%) and 570 RSV/B (83.4%) isolate sequences for analysis (see **Suppl. Table 2**) or 1176 (54.9%) of all positive cases. The distribution of sequenced samples closely mirrored that of the total positive samples, ensuring that the sequenced population was representative of the collected population. The exceptions for this were during the 2016-17 years, when RSV samples were not sequenced due to sample availability and funding issues. **Figure 1** demonstrates the effectiveness of our in-silico genotyping, showing patterns similar to the PCR classification (**Suppl. Figs. 1A and B**), with very few discordant samples noted.

**Figure 1.**
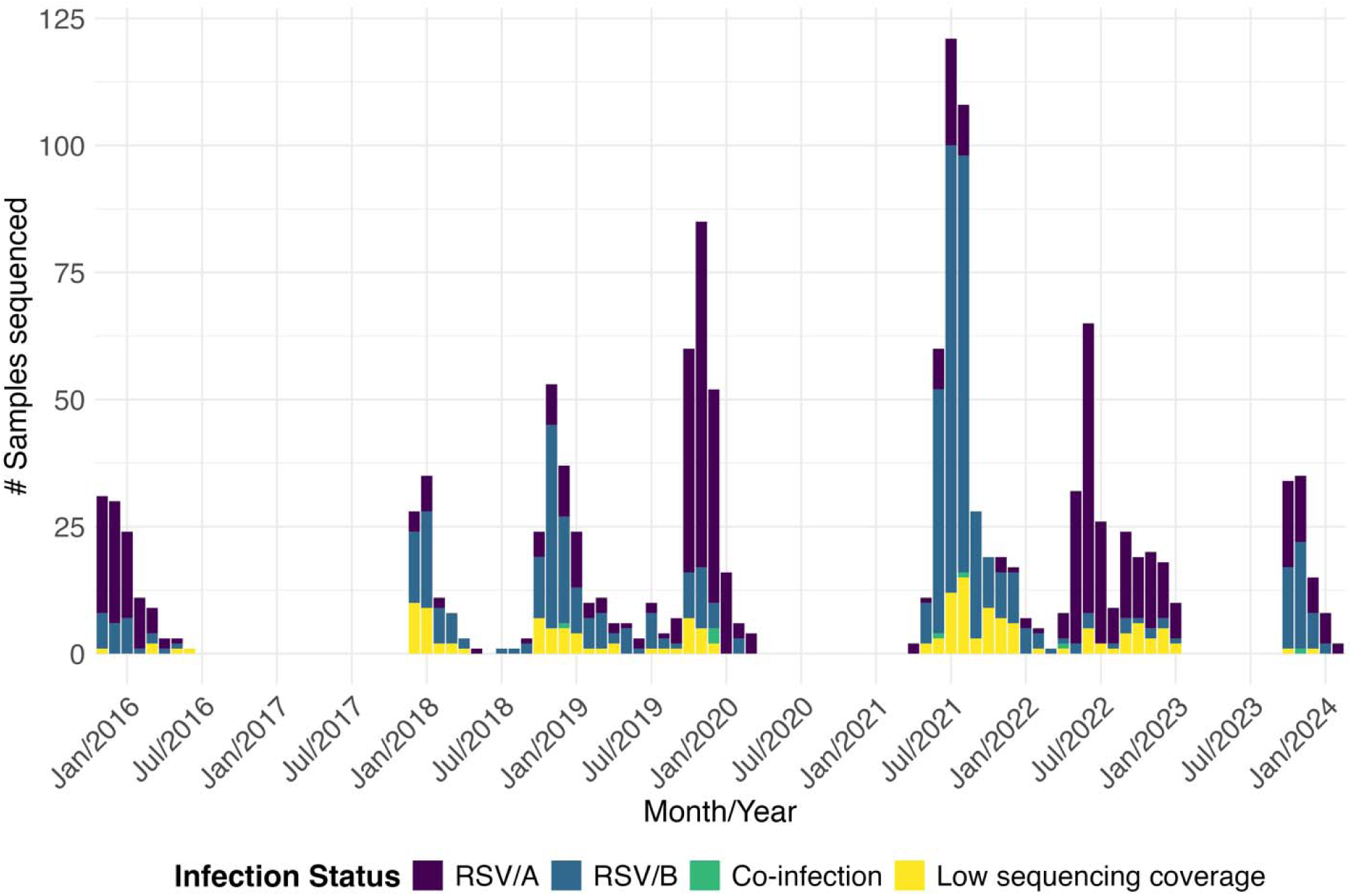
Monthly distribution of all sequenced RSV-positive samples by each subgroup and co-infection using *in-silico* classification. RSV/A, RSV/B, and co-infections of RSV/A and RSV/B are shown. Co-infection samples and low sequencing coverage samples, which did not have at least 90% of the genome covered by 20X depth, were discarded for downstream analysis.

### Phylogenetic analysis of RSV during the pre-pandemic, pandemic, and post-pandemic phases

Phylogenetic analysis revealed distinct temporal patterns in the RSV/A population structure across pre-pandemic, pandemic, and post-pandemic phases (**Figure 2**). The pre-pandemic period exhibited significant diversity, with multiple co-circulating RSV lineages, including A.D, A.D.2.2, and A.D.1. The A.D.1 lineage demonstrated remarkable persistence across all three time-phases. A notable transition occurred with the A.D.3 lineage, which was rarely observed pre-pandemic but became widespread during and after the pandemic. The A.D.5 clade diversified during the study period, with A.D.5.1 spanning all three phases and A.D.5.2 emerging during the pandemic and persisting afterward. A.D.5.3 isolates formed a distinct pre-pandemic cluster with limited pandemic-phase persistence.

**Figure 2:**
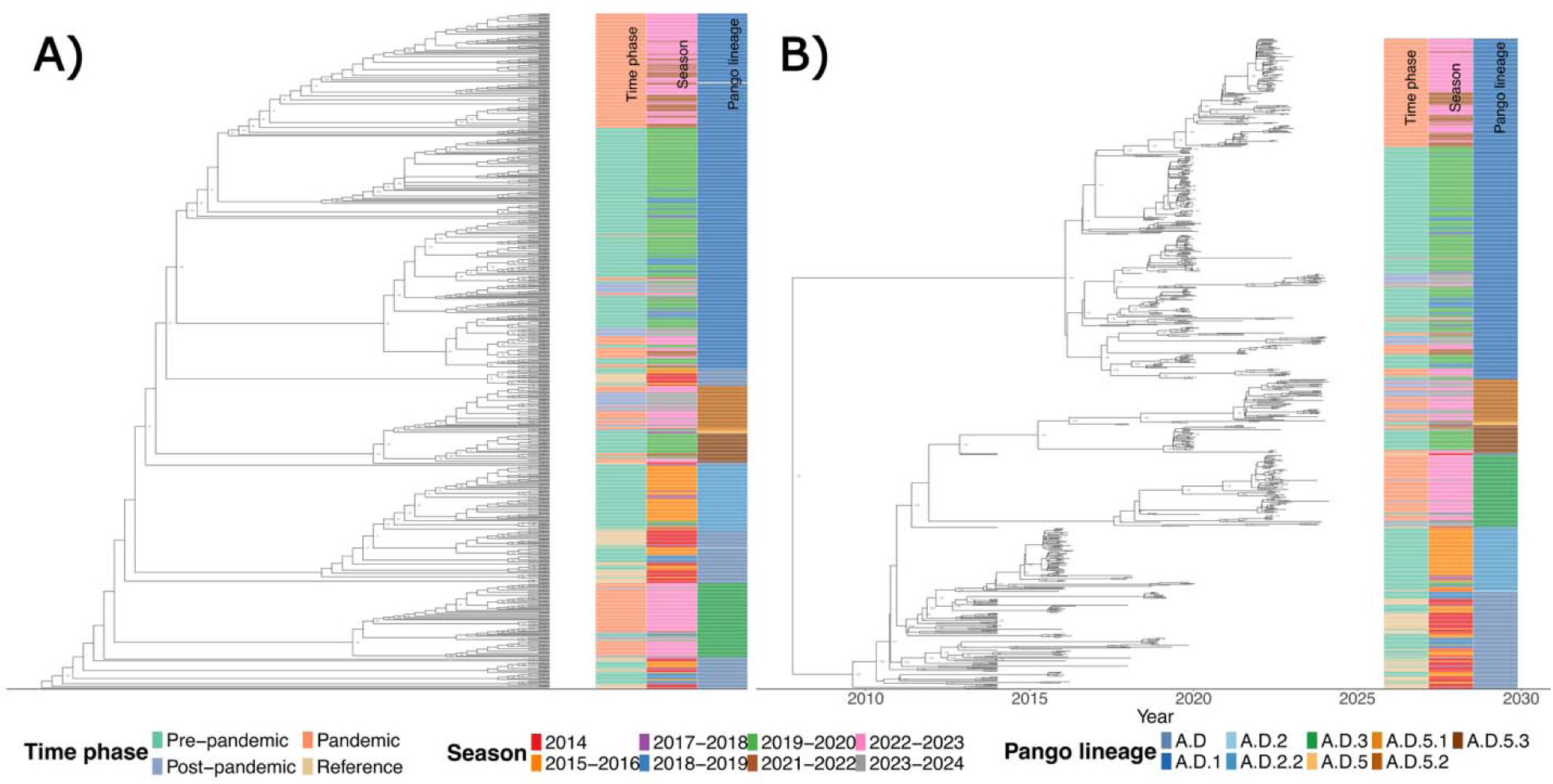
RSV/A phylogenetic analysis. (A) Cladogram showing phylogenetic clustering of RSV/A isolates from Houston, 2015–2024. (B) Time-calibrated maximum clade credibility (MCC) tree estimated by BEAST2 under a relaxed molecular clock. The x-axis represents calendar time in years. Tip annotations show time phase, season, and RSV lineage classification assigned via UShER (Goya et al., 2024). RSV/A exhibited multiple co-circulating lineages across all time phases, with A.D.1 persisting throughout the study period and A.D.3 emerging during the pandemic.

The RSV/B population (**Figure 3**) showed a more dramatic restructuring of genetic lineage during the study period. The pre-pandemic population was dominated by B.D.4.1.1 and B.D.4.1 lineages. The pandemic marked a significant transition, with most pre-existing lineages disappearing except for a few persistent B.D.4.1.1 strains. This period saw the emergence and subsequent dominance of the B.D.E.1 lineage, which maintained its prevalence into the post-pandemic phase. Some isolates assigned to the same lineage did not always form monophyletic groups in our local phylogeny; this is expected when lineage assignments are based on global reference phylogenies (via UShER), while our trees are reconstructed from local sequences only.

**Figure 3:**
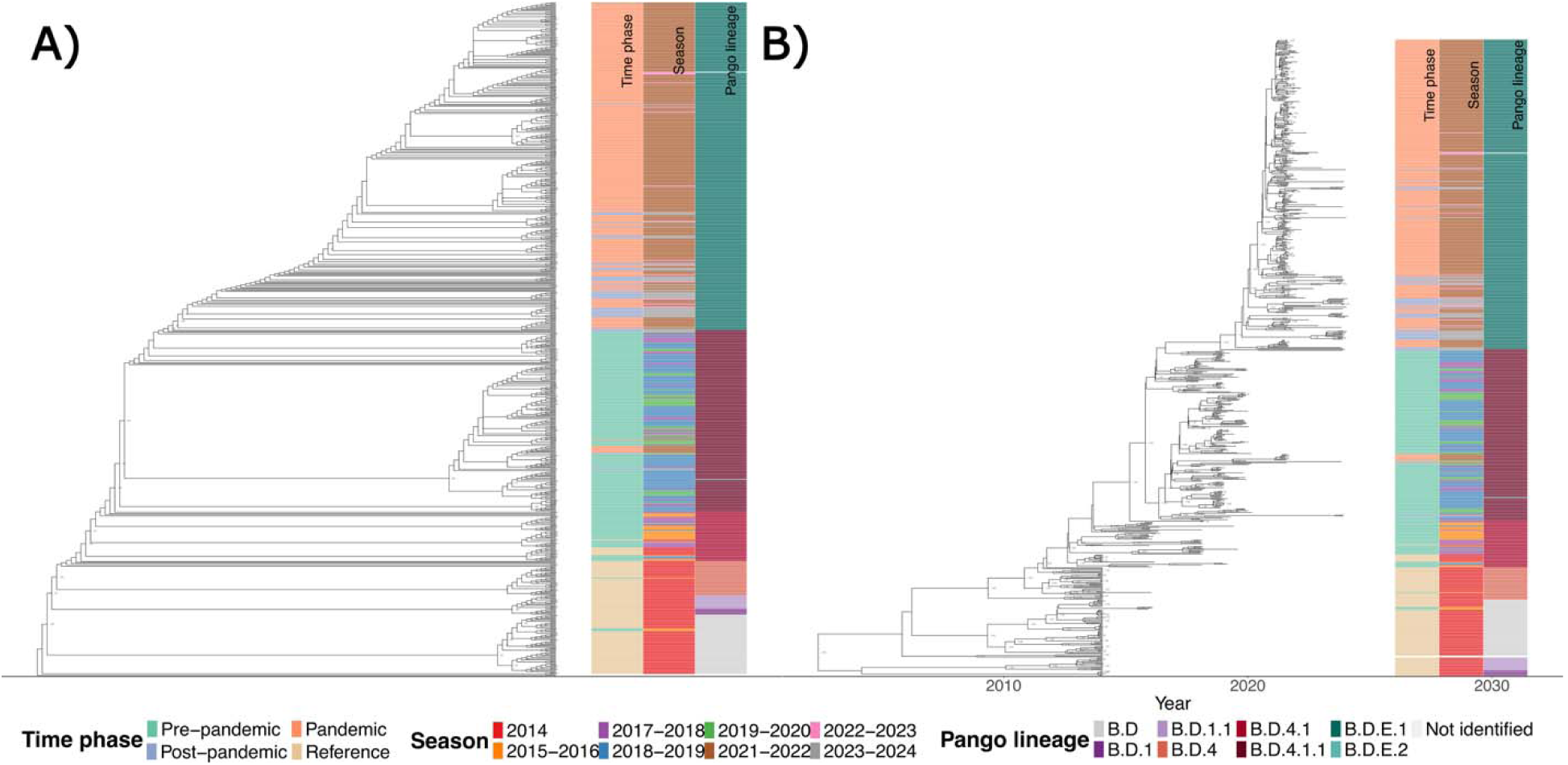
RSV/B phylogenetic analysis. (A) Cladogram showing phylogenetic clustering of RSV/B isolates from Houston, 2015–2024. (B) Time-calibrated MCC tree estimated by BEAST2. RSV/B showed a dramatic restructuring, with pre-pandemic B.D.4.1 lineages replaced by the dominant B.D.E.1 lineage during and after the pandemic.

### Molecular clock analysis and evolutionary rates

Bayesian molecular clock analysis using BEAST2 estimated mean nucleotide substitution rates of 9.37 × 10^-4^ substitutions/site/year (95% HPD: 7.99 × 10^-4^ – 1.07 × 10^-3^) for RSV/A and 9.79 × 10^-4^ substitutions/site/year (95% HPD: 8.96 × 10 – 1.06 × 10^−3^) for RSV/B. These rates are consistent with previously published estimates for RSV whole-genome sequences (Piñana et al., 2024; Tan et al., 2012). The relaxed molecular clock model was supported by substantial rate variation among branches for both subgroups (ucldStdev: 0.624 for RSV/A, 0.746 for RSV/B), indicating that a strict clock would be inappropriate for these data.

The time to the most recent common ancestor (tMRCA) of the A.D.1 RSV/A clade was estimated at approximately 2010 (95% HPD: 2008–2011), while for RSV/B it was approximately 2002 (95% HPD: 1997–2007). The tMRCA of the RSV/B lineages in our dataset was estimated at approximately 2002 (95% HPD: 1997–2007), indicating that the common ancestor of all sampled RSV/B diversity, including the dominant B.D.E.1 lineage, existed approximately two decades before the most recent samples. The time-calibrated MCC phylogenies are presented in Figures 2B and 3B.

### Genetic diversity of RSV in the pre-pandemic, pandemic, and post-pandemic phases

RSV genomic evolution during the COVID-19 pandemic showed distinct patterns between RSV/A and RSV/B subgroups. We analyzed variants in isolates collected over a 9-year period (2015-2024), spanning pre-pandemic, pandemic, and post-pandemic phases. As shown in Table 1, the number of unique variants, non-synonymous variants, and the affected samples varied by season and number of subjects for both subgroups. Analysis of total genomic variants revealed a steady accumulation in both RSV/A and RSV/B over time (**Figure 4A**). Notably, RSV/B consistently exhibited a higher baseline number of variants compared to RSV/A from the outset (**Figure 4A**, **Table 1**). Linear regression analysis demonstrated that RSV/B accumulated variants at a higher rate than RSV/A (11.57 vs. 10.65 variants per season, p=0.03) (**Figure 4B**). A list of all valid variant observations and all unique variants is available on Suppl. Tables 03 and 04, respectively.

**Figure 4.**
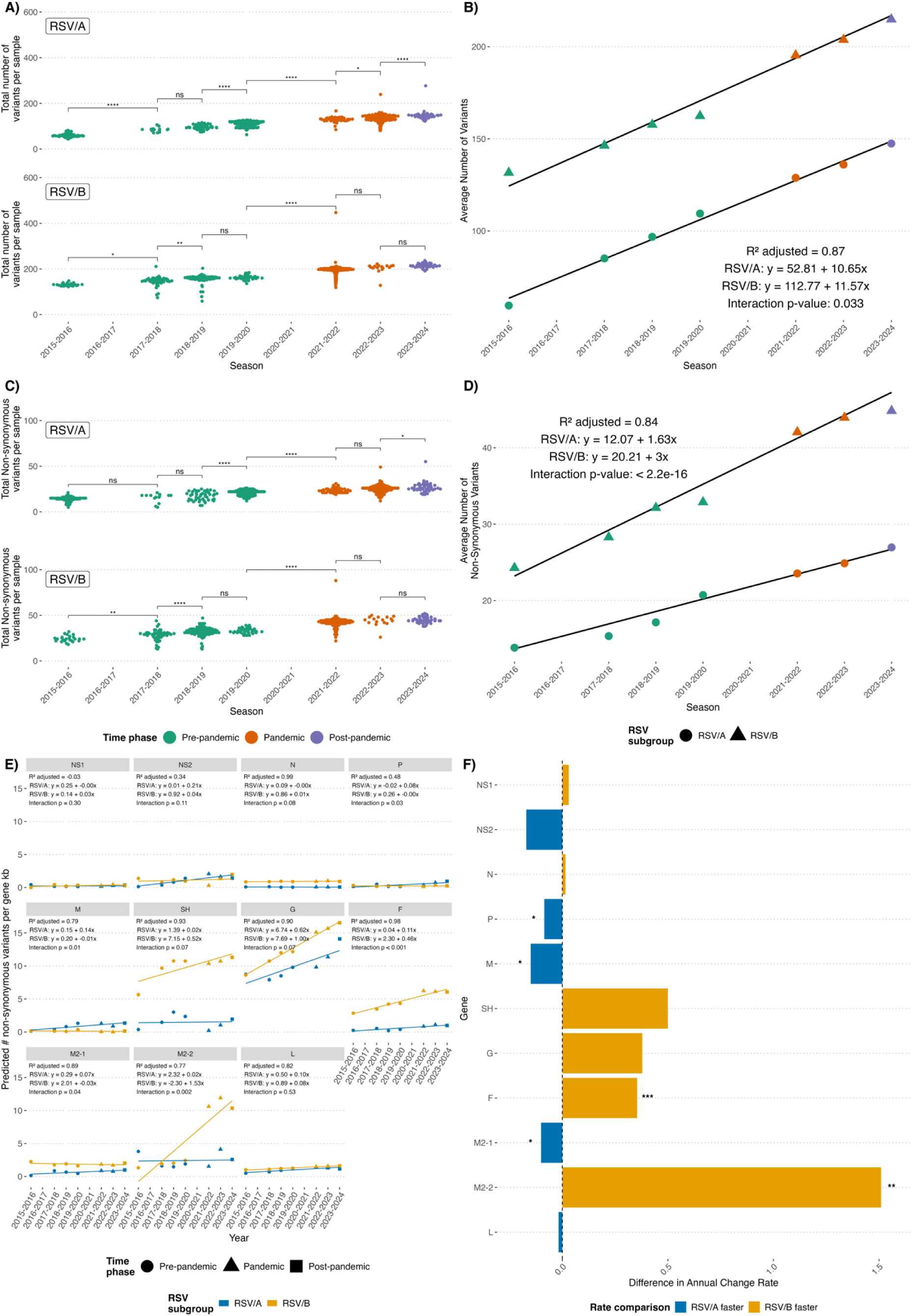
Accumulation of genomic variants in RSV/A and RSV/B subgroups across a 9-year period (2015-2024). **(A)** Total variants per isolate for RSV/A (top) and RSV/B (bottom) across consecutive seasons. Each dot represents an individual isolate, colored by time phase (pre-pandemic: green; pandemic: orange; post-pandemic: purple). Statistical significance between seasons is indicated (ns: not significant; *p<0.05; **p<0.01; ***p<0.001; ****p<0.0001; ANOVA with Tukey’s post-test). **(B)** Linear regression analysis of total variants per season for RSV/A (circles) and RSV/B (triangles). Points show the average number of variants per season for visualization purposes, while regression was performed using individual isolate data. Regression equations and adjusted R² value are shown. **(C)** Non-synonymous variants per isolate for RSV/A (top) and RSV/B (bottom) across seasons, with statistical comparisons as in panel A. **(D)** Linear regression analysis of non-synonymous variants, showing significantly different accumulation rates between RSV/A and RSV/B (p<2.2e-16). Points represent the average number of non-synonymous variants per season for visualization, while the regression was conducted on individual isolate data. **(E)** Gene-specific analysis of non-synonymous variant accumulation over time for 11 RSV genes. Each panel shows predicted variants per gene with regression statistics and interaction p-values comparing subgroups. **(F)** The difference in the annual change rate of non-synonymous variants between RSV subgroups for each gene. Positive values (orange bars) indicate faster evolution in RSV/B; negative values (blue bars) indicate faster evolution in RSV/A. Statistical significance: *p<0.05; **p<0.01; ***p<0.001.

**Table 1:** Summary of unique variants and the number of isolates per season for both RSV subgroups. A unique variant is a distinct genetic alteration identified in the RSV genome, where each unique variant may be observed across multiple isolates. P-values of the differences between RSV/A and RSV/B were calculated using One-way ANOVA with Tukey’s post-test comparing the number of non-synonymous variants for each isolate between each RSV subgroup for every season.

| Season | Time Phase | Subgroup | Number of unique variants | Number of unique non-synonymous variants | Number of subjects | p-values of the difference between RSV/A and RSV/B |
| --- | --- | --- | --- | --- | --- | --- |
| 2015-2016 | Pre-pandemic | RSV/A | 891 | 188 | 82 | 2.04E-14 |
|  |  | RSV/B | 739 | 153 | 25 |  |
| 2017-2018 | Pre-pandemic | RSV/A | 645 | 102 | 14 | 1.99E-11 |
|  |  | RSV/B | 1055 | 234 | 48 |  |
| 2018-2019 | Pre-pandemic | RSV/A | 970 | 177 | 46 | < 2.2E-16 |
|  |  | RSV/B | 1378 | 311 | 107 |  |
| 2019-2020 | Pre-pandemic | RSV/A | 1848 | 445 | 185 | 2.29E-14 |
|  |  | RSV/B | 936 | 204 | 39 |  |
| 2021-2022 | Pandemic | RSV/A | 847 | 180 | 49 | 2.01E-13 |
|  |  | RSV/B | 1708 | 446 | 289 |  |
| 2022-2023 | Pandemic | RSV/A | 1977 | 464 | 185 | < 2.2E-16 |
|  |  | RSV/B | 507 | 125 | 16 |  |
| 2023-2024 | Post-pandemic | RSV/A | 1150 | 233 | 45 | 3.99E-10 |
|  |  | RSV/B | 1132 | 246 | 46 |  |

When focusing specifically on non-synonymous variants, which alter amino acid sequences, the difference between subgroups became more pronounced (**Figure 4C**). RSV/B accumulated non-synonymous variants at nearly twice the rate of RSV/A (3.0 vs. 1.63 variants per season, p<2.2e-16) (**Figure 4D**), suggesting potentially greater adaptive evolution in the RSV/B subgroup during this period. Gene-specific analysis revealed differential patterns of non-synonymous variant accumulation across the RSV genome (**Figure 4E**). The M2-2 and F genes showed significantly higher rates of non-synonymous changes in RSV/B compared to RSV/A, while the P, M, and M2-1 genes exhibited the opposite pattern with higher rates in RSV/A (**Figure 4F**). These findings indicate gene-specific evolutionary pressures that differ between RSV subgroups, potentially reflecting their distinct adaptive strategies that emerged during the pandemic period.

### Patterns of variant prevalence for RSV/A and RSV/B

We analyzed how the evolutionary dynamics of RSV variants changed across the pre-pandemic, pandemic, and post-pandemic time phases, especially focusing on how the prevalence of genomic variants changed over time **(Figure 5)**. Variants with similar patterns of prevalence throughout the three phases clustered together, revealing distinct patterns of variant prevalence for both RSV/A and RSV/B isolates, with a clear shift in the variant landscape during the pandemic phase. RSV/A (**Figure 5A**) demonstrated variants that increased in prevalence during the pandemic followed by a decrease in post-pandemic phase (Clusters 1 and 6), variants with increased prevalence only during the post-pandemic phase (Clusters 2 and 3), variants with decreased prevalence during the pandemic with varying recovery patterns (Cluster 4 and 5), a stable subset of low-frequency variants throughout the time phases (cluster 7), and a cluster mostly of high-frequency variants that remained relatively stable during the pre-pandemic and pandemic periods (cluster 8). For RSV/B analysis (**Figure 5B**), we observed variants with dramatic prevalence increases during the pandemic (Clusters 1, 3, and 6), with variants from clusters 1 and 3 remaining stable into the post-pandemic phase, while those from cluster 6 showed a decrease in prevalence in the post-pandemic phase. Variants from cluster 2 show a prevalence decrease during the pandemic and remain low afterward. Clusters 4, 7, and 8 contain variants that were relatively stable during this study, with those from cluster 7 being of high prevalence and those from clusters 4 and 8 being of low prevalence.

**Figure 5:**
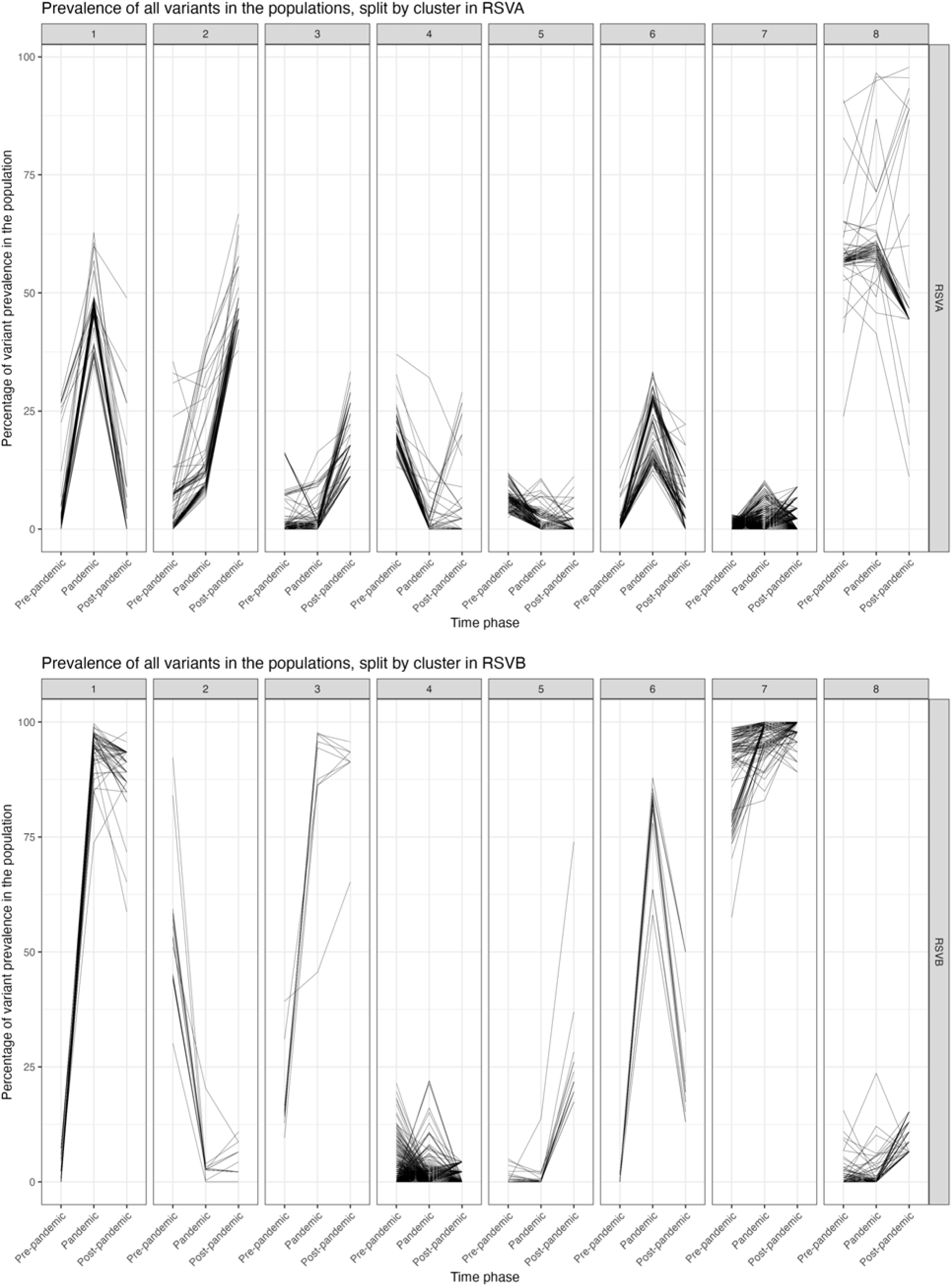
Evolutionary dynamics of RSV variants across pre-pandemic, pandemic, and post-pandemic phases. Variants are grouped into 8 clusters based on their prevalence behavior across the three time-phases. Each line in the figure represents a unique genomic variant, and the y-axis represents the scaled prevalence (population allele frequency) of that variant within each time phase. The x-axis divides the study period into three distinct phases: pre-pandemic, pandemic, and post-pandemic. Clustered prevalence patterns for RSV/A (**A**) and RSV/B (**B**) variants.

To further understand evolutionary dynamics, we quantified the association between each variant cluster (**Figure 5**) and the isolates from lineage clades identified in the phylogenetic analysis (Figures 2 and 3). This analysis revealed strong associations between specific clusters and dominant lineages, providing insights into how mutations contributed to the success or decline of different RSV subgroups during the study period (**Supplementary Table 3**). For RSV/A, variant observations from cluster 1 were predominantly associated with A.D.1 lineage, which aligns with its transient dominance during the pandemic (**Figure 5A**). This suggests that mutations in Cluster 1 may have provided a short-term fitness advantage during the pandemic but were not sustained afterward. Cluster 2 variants were primarily associated with the A.D.5.1 and A.D.5.3 lineages and became dominant during the post-pandemic period. Cluster 6 variants were strongly associated with the A.D.3 lineage (75.19%), which emerged and dominated during the pandemic, disappearing afterwards. Many of these mutations could have played a key role in the lineage’s success during the three different time phases. For RSV/B, Cluster 1 variants were overwhelmingly associated with the B.D.E.1 lineage (98.96%) that emerged during the pandemic and persisted into the post-pandemic phase (**Figure 3**). This strong association suggests that the mutations in Cluster 1 may be specific to the B.D.E.1 lineage, with some emergent mutations providing a fitness advantage and contributing to the lineage’s dominance. Cluster 2 variants were primarily linked to the B.D.4.1.1 lineage (87.57%), representing pre-pandemic lineages that were largely supplanted by B.D.E.1 during the pandemic. Cluster 4 variants displayed a more balanced distribution across B.D.4.1.1 (34.19%), B.D.E.1 (33.01%), and B.D.4.1 (18.99%) lineages, indicating a transitional phase where multiple lineages coexisted before the dominance of B.D.E.1.

An inspection to determine whether the varying cluster patterns were associated with specific genes in the RSV genome (**Suppl. Fig. 2**) revealed no clear association in RSV/A or RSV/B.

### Effect of selection pressure on the RSV genome

To assess the effects of selective pressure on each gene during the pre-pandemic, pandemic, and post-pandemic time phases, we calculated the ratio of non-synonymous to synonymous base substitutions (dN/dS) by gene for each season for both RSV/A and RSV/B (**Figure 6**). A dN/dS ratio greater than 1.0 is suggestive of positive selective pressure, A ratio equal to 1.0 suggests neutral evolution, while ratios below 1.0 are consistent with purifying selection. Our analysis revealed distinct patterns between RSV/A and RSV/B. In RSV/A, the M gene showed elevated non-synonymous substitution rates in the pre-pandemic period, stabilizing thereafter. The M2-2 gene also exhibited elevated dN/dS ratios at alternating years. Patterns consistent with purifying selection were observed for NS1, NS2, the SH (especially post-pandemic), the F, and N genes of RSV/A.

**Figure 6:**
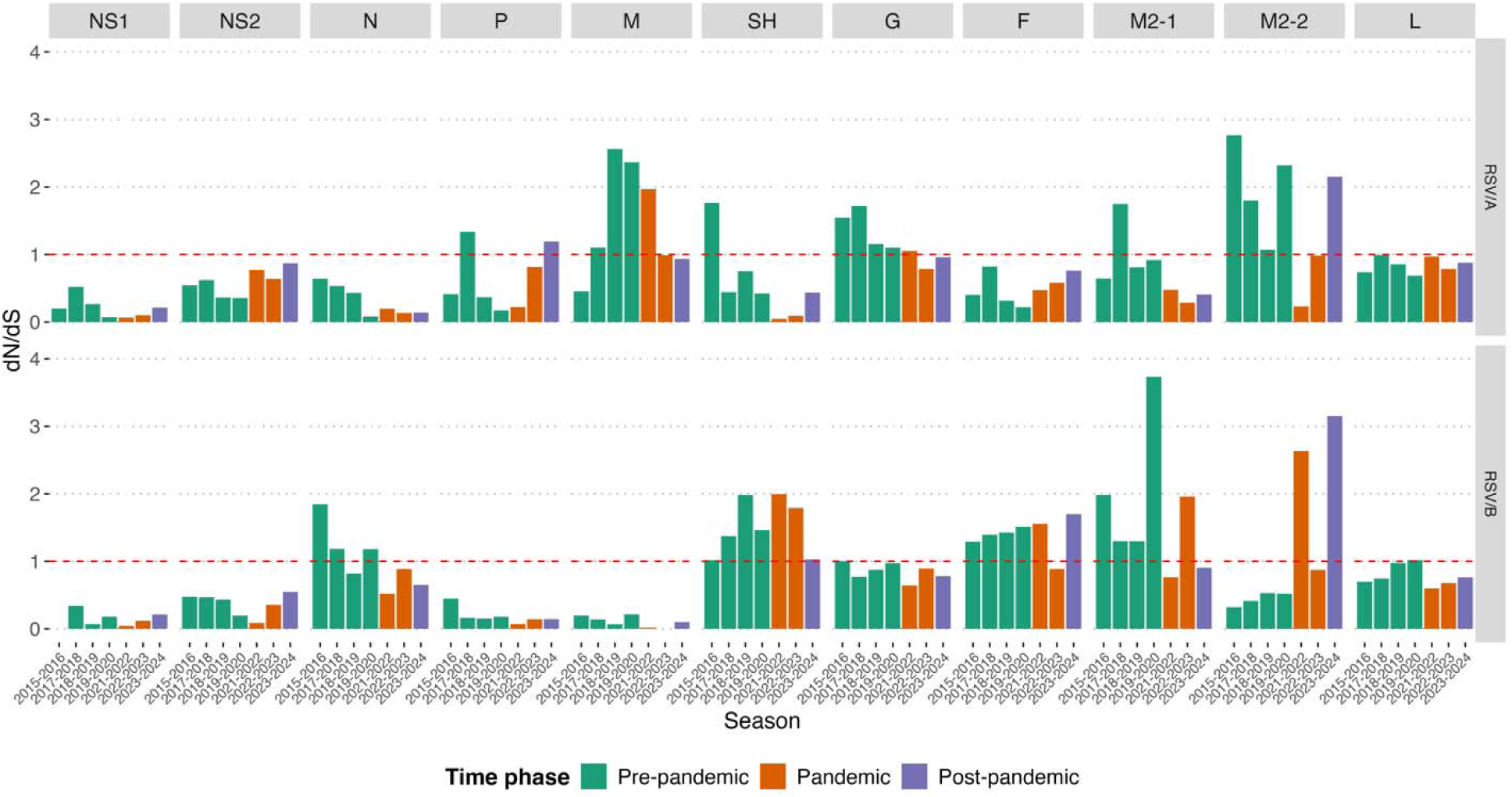
dN/dS ratios for RSV/A and RSV/B genes across seasons. The figures show the dN/dS ratios for all the genes of RSV/A (top panel) and RSV/B (bottom panel) across different seasons. Seasons are grouped into pre-pandemic (green), pandemic (orange), and post-pandemic (purple) time phases. The dN/dS ratio is plotted on the y-axis, with a dashed line at 1.0 indicating neutral selection. Values above 1.0 suggest positive selection, while values below 1.0 imply purifying selection. Each bar represents the dN/dS ratio for a specific gene in each season, allowing comparison of evolutionary pressures on different genes across time and between RSV subgroups.

Conversely, RSV/B demonstrated notably elevated dN/dS ratios for the SH and M2-1 genes during the late pre-pandemic and pandemic periods. The M2-2 gene in RSV/B showed a transition from dN/dS ratios consistent with purifying selection in the pre-pandemic period to markedly elevated ratios during the pandemic and post-pandemic phases, with a marked increase in dN/dS ratio at the onset of the pandemic. The NS1, NS2, M, and P genes in RSV/B appeared to be under purifying pressure throughout the study period.

Interestingly, the G and F genes that encode the two major surface glycoproteins inducing neutralizing antibody responses showed contrasting dN/dS patterns between the two subgroups. For RSV/A, the G gene exhibited elevated dN/dS ratios during the early pre-pandemic time phase before stabilizing near neutrality, while the F gene remained below 1.0 throughout the 9 years, consistent with purifying selection. In contrast, for RSV/B, the G gene showed near-neutral dN/dS ratios during all three time phases, while the F gene exhibited dN/dS ratios above 1.0 during all three time phases except the 2022-23 season. These findings highlight the differential evolutionary dynamics between RSV/A and RSV/B, particularly in response to the changing environment during the COVID-19 pandemic.

### Selective pressure on the M2-2 gene and its effect on transcriptional regulation

To visualize the effects of selective pressure on each gene, we plotted a simple ratio of the number of non-synonymous to synonymous variants for each isolate. As expected, the patterns closely mirrored the dN/dS ratios (**Suppl. Fig. 3**). The accumulation of non-synonymous variants differed between RSV/A and RSV/B across various genes. RSV/B isolates demonstrated a higher ratio of non-synonymous to synonymous variants in the SH and F genes compared to RSV/A in pre-pandemic, pandemic, and post-pandemic phases (**Suppl. Figure 3**).

Remarkably, though, the non-synonymous to synonymous variant ratio in the M2-2 gene remained similar for RSV/A and RSV/B in the pre-pandemic periods. However, the RSV/B non-synonymous to synonymous variant ratio was significantly higher during the pandemic and post-pandemic time phases and when compared to RSV/A (**Figure 7A**), implying a subgroup-specific evolutionary pressure or adaptation. M2-2 is a regulatory protein that helps RSV switch from transcription to RNA replication, initiating the synthesis of genomic RNA for virion assembly (Bermingham & Collins, 1999). M2-2 deletion mutants were developed as candidate RSV vaccines because of their reduced genome replication and increased viral protein production (McFarland et al, JID 2019). Given M2-2’s role in regulating transcription, we investigated whether isolates from the pandemic and post-pandemic phases (collectively termed "Pandemics+”) exhibited a lower or higher transcription ratio.

**Figure 7:**
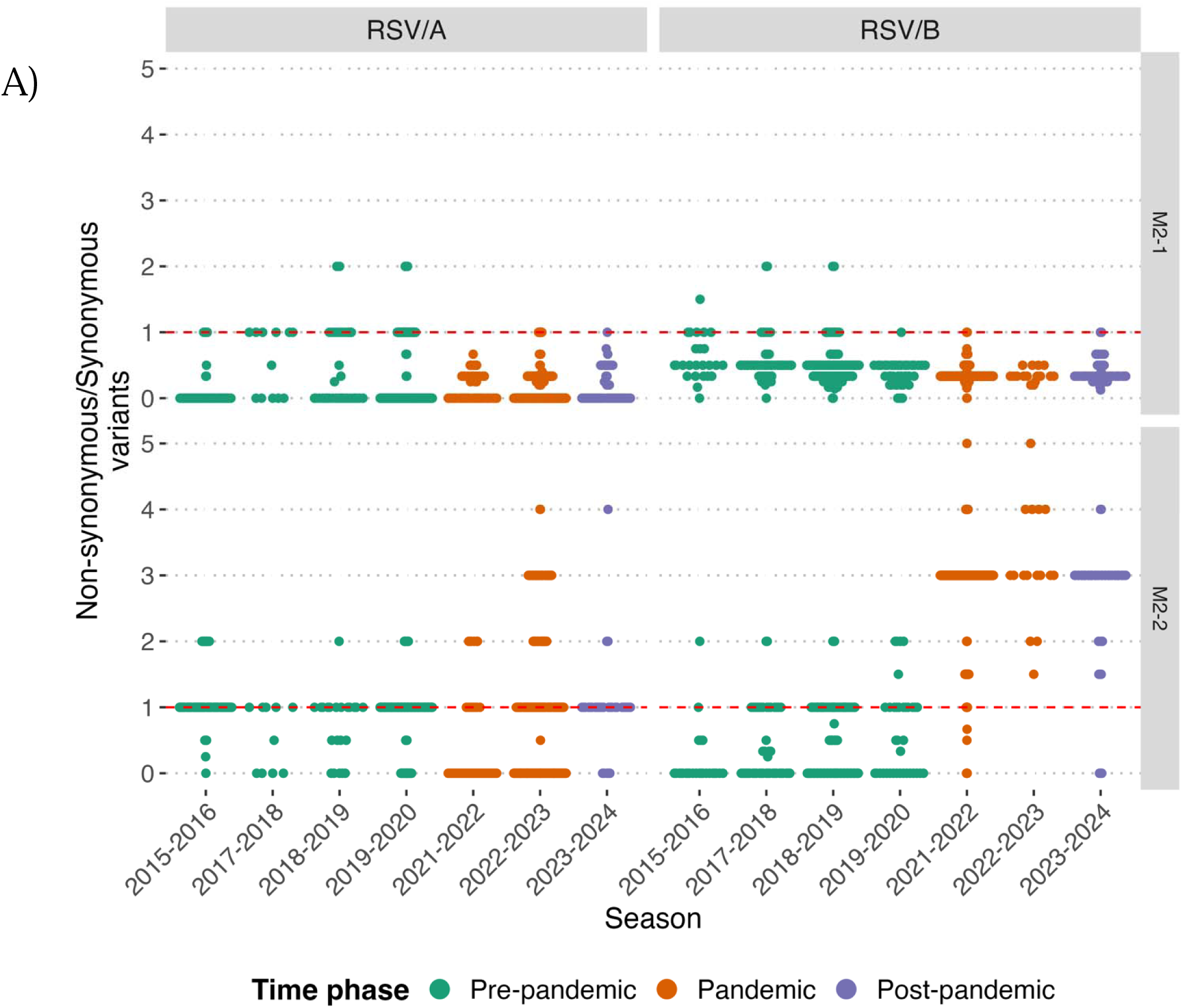
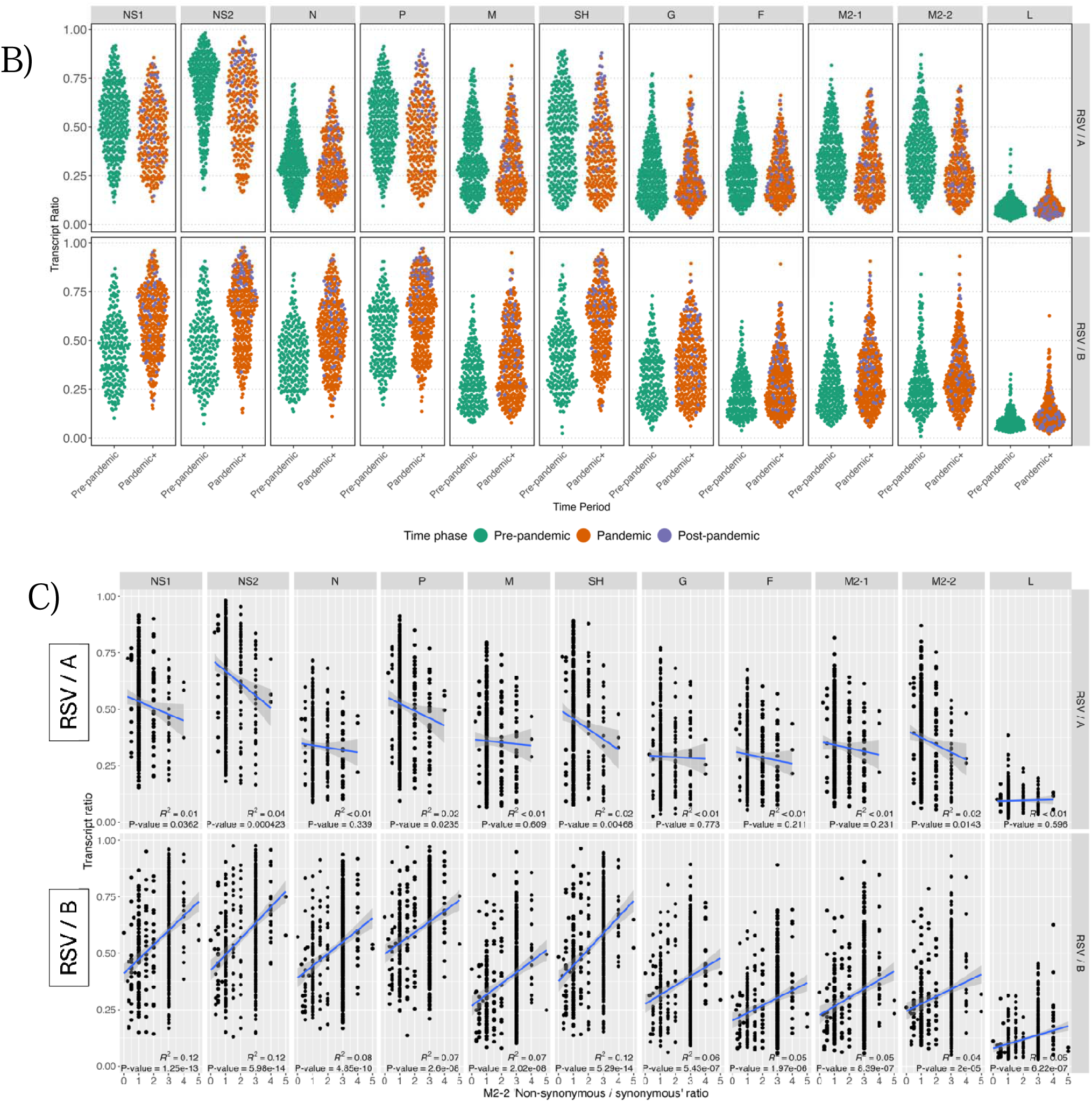
Gene-specific evolutionary dynamics and transcriptional changes in RSV during the COVID-19 pandemic. **A.** Ratio of non-synonymous to synonymous variants for the M2-1 (top) and M2-2 (bottom) genes across pre-pandemic, pandemic, and post-pandemic time phases. Data are shown separately for RSV/A (left panel) and RSV/B (right panel). Each point represents a sample, with colors indicating the time phase. **B.** Transcription ratios for RSV genes across the three time phases, derived from strand-specific sequencing libraries. The transcription ratio is calculated as the proportion of antisense strand reads (representing mRNA transcripts) relative to total reads aligned to each coding gene region. Data are presented for both RSV/A (top panel) and RSV/B (bottom panel). Each point represents a sample, color-coded by time phase as in panel A. **C.** Linear associations (R^2^) were determined between M2-2 non-synonymous/synonymous ratio and transcript ratio for RSV/A (top panel) and RSV/B (bottom panel). Each point represents a sample. R^2^ values of p <0.05 were considered statistically significant.

We previously demonstrated that probe-based capture enrichment sequencing can quantify both RSV genomes and antigenomes using strand-specific libraries (Bhamidipati et al., 2025). In a strand-specific library, reads originating from mRNA (positive-sense transcripts) align to the antisense strand of the reference genome, whereas reads derived from the negative-sense genomic RNA align to the sense strand. Using this information, we were able to calculate the ratio of transcripts to total reads to determine the proportion of transcription that occurred in each isolate. The results revealed an increase in transcript ratios for several RSV/B genes during the pandemic and post-pandemic periods (**Figure 7B**), with most (e.g., NS1, NS2, N, P, SH, G, F, M2-1, and M2-2) showing more pronounced increases. Intriguingly, a statistically significant positive linear association was observed between the non-synonymous/synonymous ratio of M2-2 and transcript/total reads ratio for all genes in RSV/B but not RSV/A. This statistically significant positive linear association strengthened the case for supporting a potential functional link between M2-2 mutations and transcriptional changes in RSV/B isolates that emerged during the pandemic (**Figure 7C**).

A deeper analysis of M2-2 variants in RSV/B revealed significant changes during the pandemic and post-pandemic periods. We identified 46 non-synonymous variants in this timeframe, with the majority (43/46) belonging to cluster 4, i.e., variants that maintained low prevalence throughout the study (**Suppl. Fig. 2B**). However, three variants showed remarkable shifts in prevalence from pre-pandemic to pandemic periods (**Figure 5C**, clusters 1 and 3, **Suppl. Fig. 2B**). Two variants (D35N and C49F) were either absent or extremely rare in the pre-pandemic period (observed in 0 and 1 out of 219 pre-pandemic samples, respectively) and became highly prevalent during the pandemic and post-pandemic phases, appearing in over 93% of the 351 isolates (cluster 1). Similarly, variant M27T increased from a presence in 14.6% of pre-pandemic isolates to 94% in the pandemic and post-pandemic phases (cluster 3). The dramatic shifts in the prevalence of these three variants and the significant positive correlation observed for RSV/B, but not RSV/A, suggest a potential functional link between one or more of these M2-2 mutations and the observed pandemic/post-pandemic increase in the transcriptional ratio across all RSV/B genes. While the strand-specific nature of our libraries supports the validity of transcript ratio measurements, these positive linear associations would benefit from experimental validation to establish a direct causal relationship. Nonetheless, these observations underscore potentially adaptive changes in RSV/B isolates that coincided with the altered epidemiological landscape during and after the COVID-19 pandemic.

### Monitoring of variants in RSV structural proteins

We next focused our analysis on variants in the G and F proteins, given their critical roles in the immune response, vaccine development, and therapeutic interventions. The F protein contains several antigenic sites (Ø, I-V). Antigenic site II is the target of the monoclonal antibody palivizumab, site V for suptavumab, site Ø for nirsevimab, and site IV for clesrovimab (Terstappen et al., 2024). Analysis of variants at antigenic sites showed more changes occurring in RSV/B and at a higher prevalence. We identified some of the previously reported site V mutations (L172Q, S173L, and K191R) that were community-acquired in RSV/B and led to suptavumab’s failure in the phase III clinical trial(Simões et al., 2021). RSV/A Site II showed several variants, including S276N with 100% prevalence in post-pandemic isolates, warranting continued monitoring. Site Ø variants I206M and Q209K/L/R were present only in RSV/B and persisted since the pre-pandemic period, also requiring ongoing surveillance (**Figure 6A).** These site Ø mutations have not yet been associated with resistance to nirsevimab (Ahani et al., 2023) .

To predict potential hot spots on the surface glycoproteins as well as the other structural and non-structural proteins, an entropy analysis was performed. The F protein of the RSV/A isolates did not show any highly variable amino acid positions within antigenic sites (**Figures 8A, 8C**), while in RSV/B (**Figures 8A, 8D**), there are highly variable amino acid positions within sites V (172, 190), Ø (206, 209), and I (389). In the G protein, we observed high variance in hypervariable regions, as expected (**Figures 8B, 8E, and 8F)** for both RSV/A and RSV/B isolates. We also analyzed the CXC3 binding motif and Central Conserved Domain (CCD) in the G protein, which is crucial for host cell binding and a potential target for future interventions. RSV/A displayed a couple of variants within the CXC3 domain, while RSV/B (**Figure 8B**) showed low-prevalence missense variants in the CCD. Analysis of the G protein consensus sequences revealed aa178 with high entropy in the CXC3 domain of RSV/A (**Figures 8E**) and aa176 in the CCD of RSV/B (**Figures 8F**).

**Figure 8:**
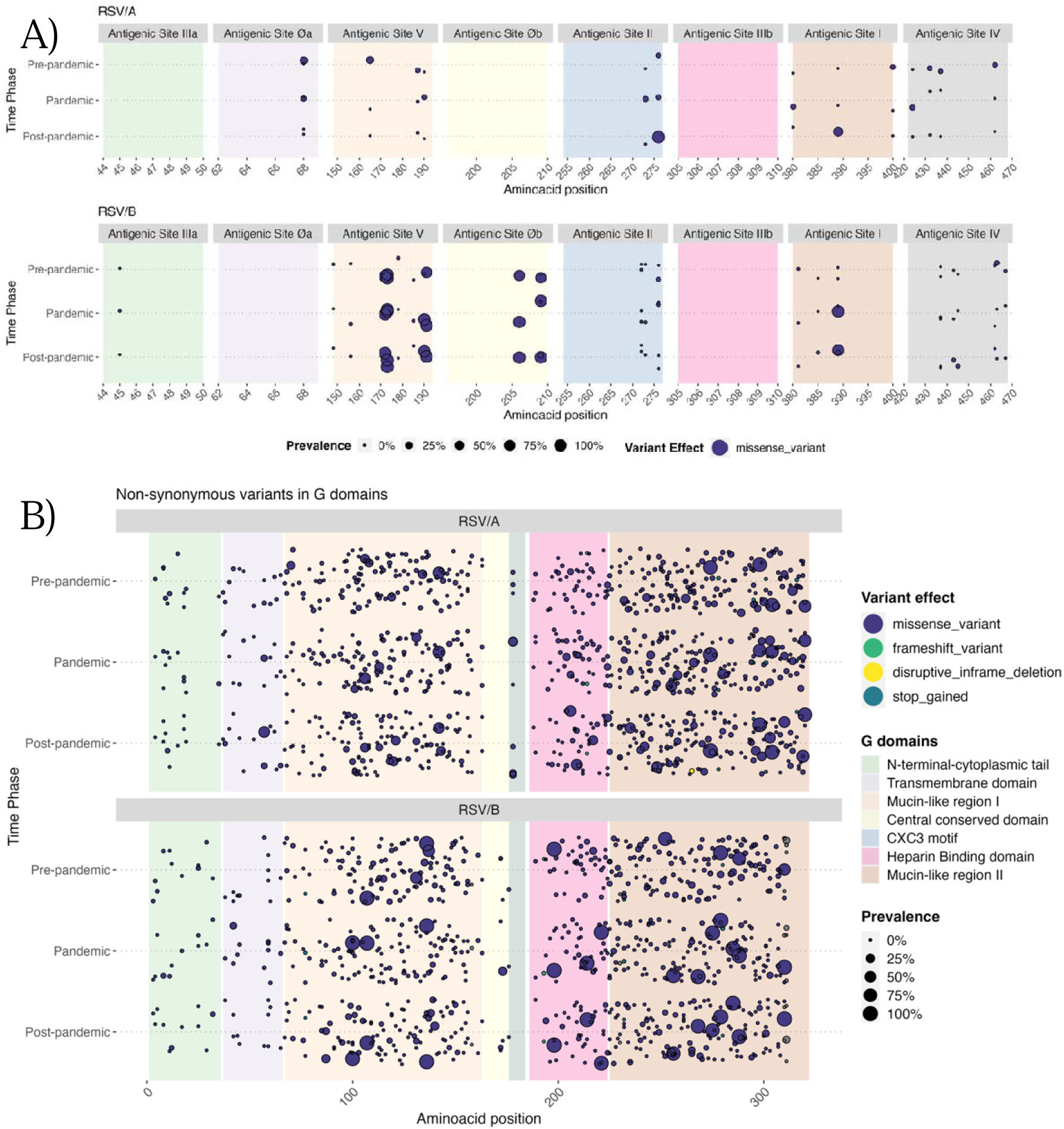
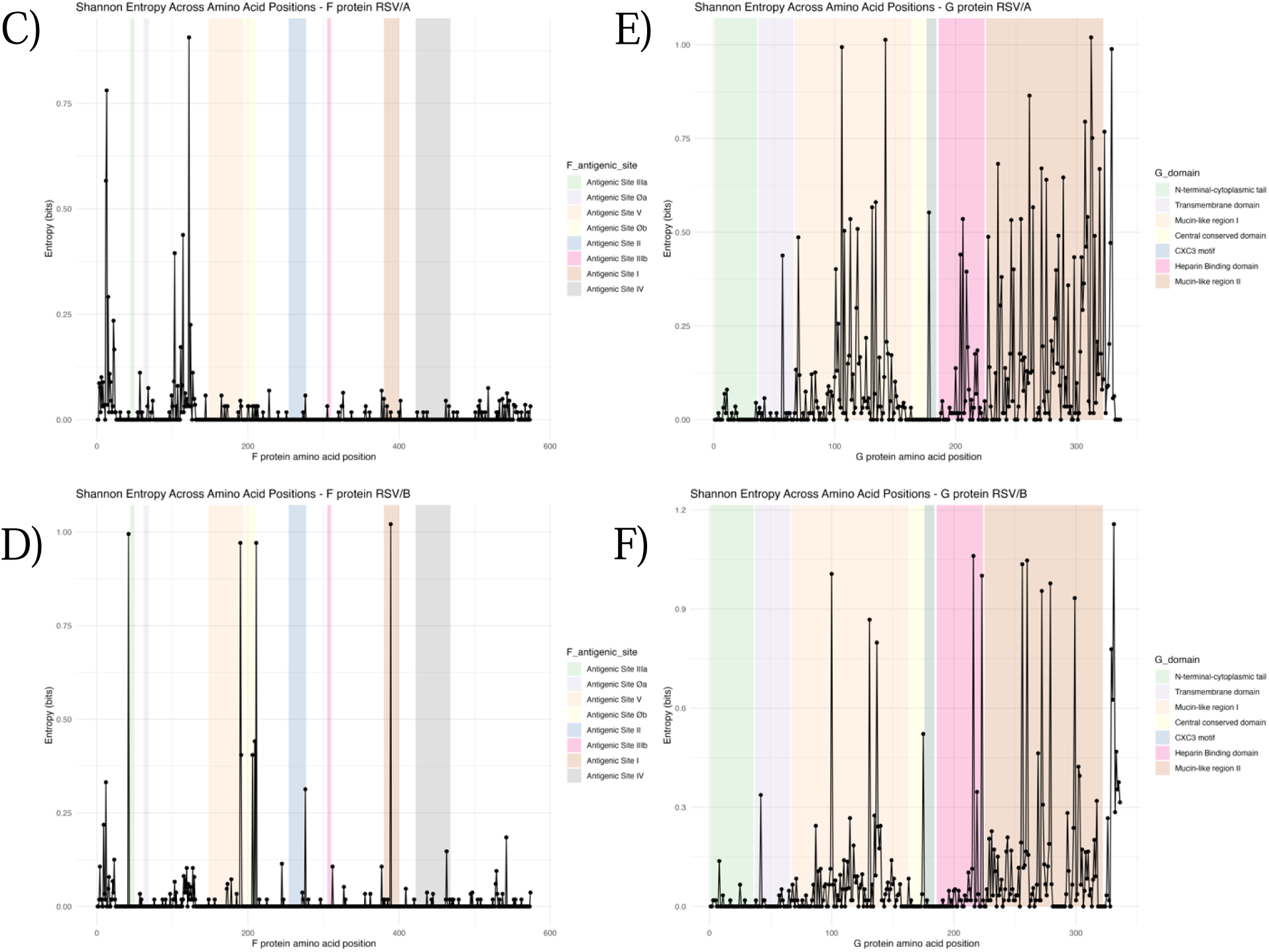
**A) and B):** All non-synonymous variants found in the antigenic sites of the F gene (A) and for the G gene (B) for RSV/A (top panel) and RSV/B (bottom panel). **C) and D)**: Shannon entropy calculated for the F protein in RSV/A (**C**) and RSV/B (**D**). Color bars represent the regions of the antigenic sites in F. **E) and F):** Shannon entropy calculated for the G protein in RSV/A (**E**) and RSV/B (**F**). Color bars represent the different G domains.

A similar analysis was performed for the rest of the genes (**Suppl. Fig. 4**). The L gene shows considerable variability in both RSV/A and RSV/B with several high entropy positions, particularly in RSV/A, suggesting ongoing evolution in this large polymerase protein. Among the non-structural and accessory proteins, M2-2 displayed notable entropy peaks in both subgroups, while the non-structural (NS1, NS2), and other structural proteins (N, P, M, SH, and M2-1) generally show lower entropy across most positions with only sporadic variability, indicating higher conservation of these genes compared to the surface glycoproteins G and F.

## Discussion

Our study provides a comprehensive analysis of the evolutionary dynamics of RSV from 1178 ARI children which is one of the largest comprehensive RSV genomic evolutionary analyses done to date. The isolates were collected over nine years, spanning pre-pandemic, pandemic, and post-pandemic time phases. By leveraging whole-genome sequencing of RSV isolates from a single geographic location, Houston, Texas, we identified key changes in RSV’s lineage dynamics, genetic diversity, and selective pressures, particularly in response to the COVID-19 pandemic, with notable differences between RSV/A and RSV/B. These findings provide crucial insights into the impact of pandemic mitigation measures on RSV circulation and the subsequent viral resurgence and may shed light on the evolutionary dynamics as vaccines and long-acting monoclonal antibody interventions are incorporated in the prevention of RSV infection among infants and older adults.

The COVID-19 pandemic led to a dramatic disruption of RSV seasonality, characterized by an initial absence of RSV circulation followed by an unusual resurgence in 2021-2023 (Dolores et al., 2022; Korsun et al., 2024). This disruption created a genetic bottleneck effect in Houston, as evidenced by a reduction in RSV genetic diversity during the pandemic period, followed by the emergence of novel lineages post-pandemic. Our phylogenetic analysis revealed that for RSV/A, the selective bottle-neck pressure led to the disappearance of several pre-pandemic lineages (A.D., A.D.2.2) while others, like A.D.3, A.D.5.2, emerged and established dominance. The persistence of certain lineages (A.D.1) across all time-phases suggests variable fitness advantages under different transmission conditions. The RSV/B population underwent an even more dramatic restructuring, characterized by near-complete replacement of pre-existing lineages by B.D.E.1, with only a small number of B.D.4.1.1 lineage showing resilience through this transition. Our findings in Houston align with global studies conducted in Australia and New Zealand, where a significant reduction was reported in RSV/A genetic diversity during the pandemic, followed by the emergence of novel lineages during the resurgence period (Eden et al., 2022; Jelley et al., 2024). Studies from other parts of the US (Rios-Guzman et al., 2024), South Africa (Sondlane et al., 2024), and Argentina (Dolores et al., 2022) have also described post-pandemic shifts in RSV lineage dominance and increased genetic diversity, supporting our observation. The emergence and dominance of B.D.E.1 in the Houston cohort mirrors trends seen across multiple regions, suggesting a global rather than local replacement event. The persistence of specific lineages, such as A.D.1 in RSV/A and B.D.E.1 in RSV/B, suggests a possible fitness advantage that warrants further investigation. Overall, the strong resurgence of both RSV subgroups post-pandemic characterized by high case numbers and increased genetic diversity, can be attributed to several factors such as susceptible population, “immune debt” in children caused by absence of RSV circulation in 2020-2021 resulting in waning immunity, potential bottleneck effect during the low-transmission period followed by the introduction of new strains from multiple geographical areas with relaxation of public health measures and resurgence in domestic and international travel, and selection pressure favoring more transmissible variants. However, restriction to local Houston sequences limits our ability to differentiate in situ evolution from external introductions and generalizability to other geographic regions with different RSV transmission dynamics.

The molecular clock analysis provided temporal context for these lineage dynamics. Estimated substitution rates for RSV/A (9.37 × 10⁻ subs/site/year) and RSV/B (9.79 × 10⁻ subs/site/year) are consistent with prior whole-genome estimates and correspond to ∼14–15 substitutions per genome per year (Piñana et al., 2024; Tan et al., 2012). The slightly higher rate and greater non-synonymous variant accumulation in RSV/B support a more dynamic evolutionary trajectory. The tMRCA for RSV/B (∼2002) is consistent with diversification of the B.D.4 and B.D.E lineages preceding pandemic-associated restructuring, whereas the tMRCA (2010) for the main RSV/A clade coincides with the introduction of the Ontario genotype with its 72 nucleotide duplication in the distal third of the G gene, that led to the rapid replacement of other RSV/A genotypes.

It would be important to note the distinction between genetic variability and effective population diversity. Although RSV/B accumulated non-synonymous variants at a higher rate, these changes occurred largely within the dominant B.D.E.1 lineage, reflecting reduced lineage diversity compared with RSV/A, which maintained multiple co-circulating clades. Thus, RSV/B showed high within-lineage mutational activity, consistent with rapid adaptation following a bottleneck, whereas RSV/A retained greater between-lineage diversity. Our variant-counting approach does not distinguish lineage-defining from convergent or transient mutations.

Our analysis of the accumulation of genetic variants across nine seasons of RSV revealed that both RSV/A and RSV/B gradually developed unique variants, as anticipated. However, the patterns of appearance and disappearance of these genomic variants differed between the two subgroups. For RSV/A, variants that emerged during the pandemic vanished in the post-pandemic period, particularly in the G, F, and L genes. In contrast, many RSV/B variants that first appeared during the pandemic persisted into the post-pandemic time phase, highlighting distinct evolutionary trajectories for the two subgroups. Furthermore, RSV/B exhibited higher dN/dS ratios across multiple genes compared to RSV/A, suggesting more dynamic evolutionary changes. Interestingly, the G genes of RSV/A but not RSV/B displayed strong positive selection, with dN/dS ratios greater than 1, indicating adaptive evolution. Conversely, other genes, such as N, P, and M, were purifying selection, which helps maintain their functional integrity. Previous studies, including a recent one by Piñana et al., have shown that RSV/B evolves at a higher rate than RSV/A and that the G gene of RSV shows the highest positive selection pressure (Lydia et al., 2013; Piñana et al., 2024; Tan et al., 2012; J.-M. Yu et al., 2021). Unique to our study, the F gene of RSV/B isolates also showed a strong selection pressure not seen for RSV/A. This observation might be highly relevant to forecasting the potential impact on the F gene of RSV/B isolates as the monoclonal antibody, nirsevimab and clesrovimab, coverage increases among infants. Our dN/dS analysis provides gene-level evidence of differential selective pressures but does not resolve selection at individual codons. Statistical methods may provide a more robust identification of adaptively evolving sites and distinction between episodic and pervasive selection. Moreover, we did not sequence samples from the 2016–2017 season, and we had variable sample sizes across seasons, which could have affected phylogenetic continuity and prevalence of variants

One of the most striking findings in our study was a significant increase in non-synonymous mutations (D35N, 49F, and M27T) observed in the M2-2 gene of RSV/B, particularly during the pandemic and post-pandemic phases. High sequence variability in the M2-2 has been reported previously, although in RSV/A and not in RSV/B (Tan et al., 2012) RSV virions lacking the M2-2 gene are less efficient in cell culture, resulting in lower viral loads and a marked increase in viral transcripts (Bermingham & Collins, 1999). Intriguingly, our study found that the mutations in M2-2 were highly correlated with increased transcript ratios, suggesting a potential adaptive mechanism to enhance viral replication. However, this phenomenon was not observed in RSV/A. Previous reports indicated that the loss of the M2-2 gene leads to increased expression of structural proteins G and F, resulting in greater immunogenicity despite lower replication. This finding contributed to the development and testing of live attenuated RSV vaccines that lack the M2-2 gene (Karron et al., 2015; McFarland et al., 2019). In our study, the three mutations in M2-2 resulted in a significant increase in G, NS1, and NS2 transcripts, suggesting immunomodulatory effects. Overall, considering the role of M2-2 in the switch between transcription and viral replication, we hypothesize that these mutations in RSV/B can confer a fitness advantage. Future studies should explore the functional consequences of these mutations in vitro and in vivo to assess their impact on viral replication dynamics, immune evasion, and disease severity.

A detailed analysis of antigenic site mutations in the F protein revealed significant alterations in sites V, Ø, and I, as well as II, which were consistent with previous work on RSV immune evasion mechanisms. Notably, community-acquired mutations at site V (L172Q, S173L, and K191R) have been previously reported in association with suptavumab resistance (Simões et al., 2021). Mutations I206M and Q209K/L/R in Site Ø, the target of nirsevimab antibody, were present only in RSV/B and were persistent since the pre-pandemic period. These mutations have emerged in the community without selective pressure of the antibody and have not resulted in resistance to the monoclonal (Ahani et al., 2023; Hammitt et al., 2022; Piñana et al., 2024). It is important to note that majority of the RSV isolates analyzed in our study represent pre-intervention baselines (2015-2023), Consequently, selective pressure from nirsevimab, clesrovimab, and maternal RSV vaccination was minimal, These changes underscore the potential for ongoing antigenic drift in RSV and reinforce the need for continued genomic surveillance to monitor the impact of vaccines and monoclonal antibody interventions on viral evolution.

It is important to acknowledge that not all observed changes necessarily represent adaptive evolution. Some patterns of lineage persistence or extinction following the COVID-19 pandemic may reflect founder effects, transmission bottlenecks, or stochastic drift rather than selection for increased fitness. The reduced population size of RSV during the pandemic period would have amplified the role of genetic drift, potentially allowing fixation of neutral or mildly deleterious variants that would otherwise have been purged by purifying selection. Disentangling the relative contributions of selection versus drift would require larger, multi-site datasets and functional validation of candidate adaptive mutations.

## Conclusions

Our study reveals the complex evolutionary dynamics of RSV in Houston across pre-pandemic, pandemic, and post-pandemic periods. The COVID-19 pandemic significantly disrupted RSV seasonality, eliminating typical patterns before triggering off-season surges and new lineage emergence. RSV/A and RSV/B showed distinct evolutionary paths, with RSV/B displaying faster accumulation of non-synonymous variants within a reduced number of co-circulating lineages. Specific M2-2 mutations that increased during and after the pandemic correlated with enhanced transcriptional activity, suggesting adaptations in viral replication or immune evasion. F protein mutations, particularly in RSV/B, indicate ongoing immune escape potential that may affect monoclonal antibody and vaccine efficacy.

These results highlight RSV’s evolutionary responsiveness to changing epidemiological conditions and introduction of new interventions. Continued genomic surveillance remains essential for monitoring new variants and their impact on disease severity, transmission, and prevention strategies. As RSV vaccines and monoclonal antibodies become more widely used, understanding these evolutionary pressures will be crucial for optimizing interventions and reducing RSV burden in vulnerable populations.

## Data availability

The datasets generated and or analyzed are available in NCBI GenBank and SRA, under the accession numbers PRJNA1195144. The consensus genomes of RSV from this study are being deposited in GenBank database (accession numbers pending)

## Ethics approval and consent

Institutional review board approvals were obtained at the CDC and Baylor College of Medicine. Informed written consent was obtained from parents and/or legal guardians of eligible children for participation in standardized parent and/or guardian interviews, medical chart review, and collection and testing of respiratory samples.

## Competing interests

FJS has research funding from Illumina, Pacbio, Genentech and Oxford Nanopore PAP has funding from Mapp Biologics, Merck, Breakwater Therapeutics, and Blue Lake Biotechnology, and does ad hoc consulting for Merck, Sanofi Pasteur, Pfizer, Enanta, Gilead.

## Funding

This work was supported by NIH grant U19AI144297 and the US Centers for Disease Control and Prevention (cooperative agreement RFA-IP-16-004). Disclaimer: The findings and conclusions in this report are those of the authors and do not necessarily represent the official position of the Centers for Disease Control and Prevention.

## Acknowledgments

We thank all members of the New Vaccine Surveillance Network as well as the children and parents who participated in the study.

## Notes

### Competing Interest Statement

FJS has research funding from Illumina, Pacbio, Genentech and Oxford Nanopore. FJS has research funding from Illumina, Pacbio, Genentech and Oxford Nanopore. PAP has funding from Mapp Biologics, Merck, Breakwater Therapeutics, and Blue Lake Biotechnology, and does ad hoc consulting for Merck, Sanofi Pasteur, Pfizer, Enanta, Gilead.

### Summary of Updates

With additional analysis and text to abstract, methods, results, and discussion.

## References

1. Adams, O., Bonzel, L., Kovacevic, A., Mayatepek, E., Hoehn, T., & Vogel, M. (2010). Palivizumab-Resistant Human Respiratory Syncytial Virus Infection in Infancy. Clinical Infectious Diseases, 51(2), 185–188. 10.1086/653534

2. Ahani, B., Tuffy, K. M., Aksyuk, A. A., Wilkins, D., Abram, M. E., Dagan, R., Domachowske, J. B., Guest, J. D., Ji, H., Kushnir, A., Leach, A., Madhi, S. A., Mankad, V. S., Simões, E. A. F., Sparklin, B., Speer, S. D., Stanley, A. M., Tabor, D. E., Hamrén, U. W., … Villafana, T. (2023). Molecular and phenotypic characteristics of RSV infections in infants during two nirsevimab randomized clinical trials. Nature Communications, 14(1), 4347. 10.1038/s41467-023-40057-8

3. Aiken, L. (1991). Multiple regression: Testing and interpreting interactions. https://books.google.com/books?hl=en&lr=&id=LcWLUyXcmnkC&oi=fnd&pg=PP11&ots=fqilXhUU2d&sig=BItmcpX1GJ5A20maOA51UaOtfRc

4. Ajami, N. J., Wong, M. C., Ross, M. C., Lloyd, R. E., & Petrosino, J. F. (2018). Maximal viral information recovery from sequence data using VirMAP. Nature Communications, 9(1), 3205. 10.1038/s41467-018-05658-8

5. Alberto, P., G, I. M., M, L. J., Dong-Gun, L., Isabel, L.-R., Federico, M.-T., F, S. T., N, van Z.-S. R., Laura, C., Nancy, D., Nathalie, de S., Laurence, F., Marie-Pierre, D., Marie, V. der W., Lusine, K., & Veronica, H. (2023). Respiratory Syncytial Virus Prefusion F Protein Vaccine in Older Adults. New England Journal of Medicine, 388(7), 595–608. 10.1056/NEJMoa2209604

6. Anderson, L. J., Hierholzer, J. C., Tsou, C., Hendry, R. M., Fernie, B. F., Stone, Y., & McIntosh, K. (1985). Antigenic characterization of respiratory syncytial virus strains with monoclonal antibodies. The Journal of Infectious Diseases, 151(4), 626–633. http://www.ncbi.nlm.nih.gov/pubmed/2579169

7. Avadhanula, V., Agustinho, D. P., Menon, V. K., Chemaly, R. F., Shah, D. P., Qin, X., Surathu, A., Doddapaneni, H., Muzny, D. M., Metcalf, G. A., Cregeen, S. J., Gibbs, R. A., Petrosino, J. F., Sedlazeck, F. J., & Piedra, P. A. (2024). Inter and intra-host diversity of RSV in hematopoietic stem cell transplant adults with normal and delayed viral clearance. Virus Evolution, 10(1), vead086. 10.1093/ve/vead086

8. Avadhanula, V., Chemaly, R. F., Shah, D. P., Ghantoji, S. S., Azzi, J. M., Aideyan, L. O., Mei, M., & Piedra, P. A. (2015). Infection with novel respiratory syncytial virus genotype Ontario (ON1) in adult hematopoietic cell transplant recipients, Texas, 2011-2013. Journal of Infectious Diseases, *211*(4). 10.1093/infdis/jiu473

9. Beate, K., A, M. S., Iona, M., F, S. E. A., A, P. B., Conrado, L., Jeffrey, B., Gonzalo, P. M., David, R., Emma, S., Julia, G., Hasra, S., James, B., Philip, Z., L, B. S., Merlin, F., Tyler, A., Nicole, P., A, V. H. M., … Alejandra, G. (2023). Bivalent Prefusion F Vaccine in Pregnancy to Prevent RSV Illness in Infants. New England Journal of Medicine, 388(16), 1451–1464. 10.1056/NEJMoa2216480

10. Bermingham, A., & Collins, P. L. (1999). The M2–2 protein of human respiratory syncytial virus is a regulatory factor involved in the balance between RNA replication and transcription. Proceedings of the National Academy of Sciences of the United States of America, 96(20), 11259. 10.1073/PNAS.96.20.11259

11. Bhamidipati, S. V, Surathu, A., Chao, H., Agustinho, D. P., Xiang, Q., Kottapalli, K., Santhanam, A., Momin, Z., Walker, K., Menon, V. K., Weissenberger, G., Emerick, N., Mahjabeen, F., Meng, Q., Hu, J., Sucgang, R., Henke, D., Sedlazeck, F. J., Khan, Z. M., … Doddapaneni, H. (2025). Complete genomic characterization of global pathogens respiratory syntical virus and human norovirus using probe based capture enrichment. Scientific Reports, 15(1), 20526. 10.1038/s41598-025-03398-6

12. Bolger, A. M., Lohse, M., & Usadel, B. (2014). Trimmomatic: a flexible trimmer for Illumina sequence data. *Bioinformatics (Oxford*, England), 30(15), 2114–2120. 10.1093/bioinformatics/btu170

13. Bouckaert, R., Vaughan, T. G., Barido-Sottani, J., Duchêne, S., Fourment, M., Gavryushkina, A., Heled, J., Jones, G., Kühnert, D., De Maio, N., Matschiner, M., Mendes, F. K., Müller, N. F., Ogilvie, H. A., Du Plessis, L., Popinga, A., Rambaut, A., Rasmussen, D., Siveroni, I., … Drummond, A. J. (2019). BEAST 2.5: An advanced software platform for Bayesian evolutionary analysis. PLOS Computational Biology, 15(4), e1006650. 10.1371/JOURNAL.PCBI.1006650

14. CDC. (2025). NVSN. https://www.cdc.gov/nvsn/php/about/index.html

15. Cingolani, P., Platts, A., Wang, L. L., Coon, M., Nguyen, T., Wang, L., Land, S. J., Lu, X., & Ruden, D. M. (2012). A program for annotating and predicting the effects of single nucleotide polymorphisms, SnpEff: SNPs in the genome of Drosophila melanogaster strain w1118; iso-2; iso-3. *Fly*, *6*(2), 80–92. 10.4161/fly.19695

16. Collins, P. L., & Melero, J. A. (2011). Progress in understanding and controlling respiratory syncytial virus: Still crazy after all these years. Virus Research, 162(1–2), 80–99. 10.1016/j.virusres.2011.09.020

17. Danecek, P., Bonfield, J. K., Liddle, J., Marshall, J., Ohan, V., Pollard, M. O., Whitwham, A., Keane, T., McCarthy, S. A., Davies, R. M., & Li, H. (2021). Twelve years of SAMtools and BCFtools. GigaScience, 10(2). 10.1093/gigascience/giab008

18. Dolores, A., Stephanie, G., Mercedes S, N. J., Érica, G., Mistchenko, A. S., & Mariana, V. (2022). RSV reemergence in Argentina since the SARS-CoV-2 pandemic. Journal of Clinical Virology, 149, 105126. 10.1016/j.jcv.2022.105126

19. Drummond, A. J., Ho, S. Y. W., Phillips, M. J., & Rambaut, A. (2006). Relaxed Phylogenetics and Dating with Confidence. PLOS Biology, 4(5), e88. 10.1371/JOURNAL.PBIO.0040088

20. E, W. E., Gonzalo, P. M., M, Z. A., R, F. A., Qin, J., Michael, P., P, P. F., Conrado, L., A, D. P., Kumar, I., Mika, R., Yasushi, F., Nazreen, H., J, B. L., Jose, C., Elliot, D., Giselle, C. V., Marinela, I., Daniel, E., … Beate, S.-T. (2023). Efficacy and Safety of a Bivalent RSV Prefusion F Vaccine in Older Adults. New England Journal of Medicine, 388(16), 1465–1477. 10.1056/NEJMoa2213836

21. Eden, J.-S., Sikazwe, C., Xie, R., Deng, Y.-M., Sullivan, S. G., Michie, A., Levy, A., Cutmore, E., Blyth, C. C., Britton, P. N., Crawford, N., Dong, X., Dwyer, D. E., Edwards, K. M., Horsburgh, B. A., Foley, D., Kennedy, K., Minney-Smith, C., Speers, D., … group, the A. R. S. V. study. (2022). Off-season RSV epidemics in Australia after easing of COVID-19 restrictions. Nature Communications, *13*(1), 2884. 10.1038/s41467-022-30485-3

22. Edgar, R. C. (2004). MUSCLE: multiple sequence alignment with high accuracy and high throughput. Nucleic Acids Research, 32(5), 1792–1797. 10.1093/nar/gkh340

23. Eleanor, W., Jaya, G., H, B. A., A, D. P., Gonzalo, P.-M., Khalequ, Z., Jorge, M., A, D. C. J., Mugen, U., Mika, R., Lina, P.-B., R, F. A., E, W. E., Rakesh, D., Lauren, W., Jiejun, D., Parinaz, G., Archana, K., Lan, L., … L, C. G. (2023). Efficacy and Safety of an mRNA-Based RSV PreF Vaccine in Older Adults. New England Journal of Medicine, 389(24), 2233–2244. 10.1056/NEJMoa2307079

24. Falsey, A., Hennessey, P., Formica, M., Cox, C., & Walsh, E. (2005). Respiratory Syncytial Virus Infection in Elderly and High-Risk Adults. The New England Journal of Medicine, 352(17). 10.1056/NEJMOA043951

25. Glezen, W. P., Taber, L. H., Frank, A. L., & Kasel, J. A. (1986). Risk of primary infection and reinfection with respiratory syncytial virus. American Journal of Diseases of Children (1960), *140*(6), 543–546. http://www.ncbi.nlm.nih.gov/pubmed/3706232

26. Goya, S., Ruis, C., Neher, R. A., Meijer, A., Aziz, A., Hinrichs, A. S., von Gottberg, A., Roemer, C., Amoako, D. G., Acuña, D., McBroome, J., Otieno, J. R., Bhiman, J. N., Everatt, J., Muñoz-Escalante, J. C., Ramaekers, K., Duggan, K., Presser, L. D., Urbanska, L., … Viegas, M. (2024). Standardized Phylogenetic Classification of Human Respiratory Syncytial Virus below the Subgroup Level. Emerging Infectious Diseases, 30(8), 1631–1641. 10.3201/eid3008.240209

27. Grubaugh, N. D., Gangavarapu, K., Quick, J., Matteson, N. L., De Jesus, J. G., Main, B. J., Tan, A. L., Paul, L. M., Brackney, D. E., Grewal, S., Gurfield, N., Van Rompay, K. K. A., Isern, S., Michael, S. F., Coffey, L. L., Loman, N. J., & Andersen, K. G. (2019). An amplicon-based sequencing framework for accurately measuring intrahost virus diversity using PrimalSeq and iVar. Genome Biology, 20(1), 8. 10.1186/s13059-018-1618-7

28. Hamid, S., Winn, A., Parikh, R., Jones, J. M., McMorrow, M., Prill, M. M., Silk, B. J., Scobie, H. M., & Hall, A. J. (2023). Seasonality of Respiratory Syncytial Virus - United States, 2017-2023. MMWR. Morbidity and Mortality Weekly Report, *72*(14), 355–361. 10.15585/mmwr.mm7214a1

29. Hammitt, L. L., Dagan, R., Yuan, Y., Baca Cots, M., Bosheva, M., Madhi, S. A., Muller, W. J., Zar, H. J., Brooks, D., Grenham, A., Wählby Hamrén, U., Mankad, V. S., Ren, P., Takas, T., Abram, M. E., Leach, A., Griffin, M. P., & Villafana, T. (2022). Nirsevimab for Prevention of RSV in Healthy Late-Preterm and Term Infants. New England Journal of Medicine, 386(9), 837–846. 10.1056/NEJMoa2110275

30. Hatter, L., Eathorne, A., Hills, T., Bruce, P., & Beasley, R. (2021). Respiratory syncytial virus: paying the immunity debt with interest. The Lancet Child & Adolescent Health, 5(12), e44–e45. 10.1016/S2352-4642(21)00333-3

31. Hause, A. M., Henke, D. M., Avadhanula, V., Shaw, C. A., Tapia, L. I., & Piedra, P. A. (2017). Sequence variability of the respiratory syncytial virus (RSV) fusion gene among contemporary and historical genotypes of RSV/A and RSV/B. PLoS ONE, 12(4). 10.1371/journal.pone.0175792

32. Jelley, L., Douglas, J., O’Neill, M., Berquist, K., Claasen, A., Wang, J., Utekar, S., Johnston, H., Bocacao, J., Allais, M., de Ligt, J., Tan, C. E., Seeds, R., Wood, T., Aminisani, N., Jennings, T., Welch, D., Turner, N., McIntyre, P., … Geoghegan, J. L. (2024). Spatial and temporal transmission dynamics of respiratory syncytial virus in New Zealand before and after the COVID-19 pandemic. Nature Communications 2024 15:1, 15(1), 1–11. 10.1038/s41467-024-53998-5

33. Karron, R. A., Luongo, C., Thumar, B., Loehr, K. M., Englund, J. A., Collins, P. L., & Buchholz, U. J. (2015). A gene deletion that up-regulates viral gene expression yields an attenuated RSV vaccine with improved antibody responses in children. Science Translational Medicine, 7(312). 10.1126/SCITRANSLMED.AAC8463/SUPPL_FILE/7-312RA175_SM.PDF

34. Korsun, N., Trifonova, I., Madzharova, I., Alexiev, I., Uzunova, I., Ivanov, I., Velikov, P., Tcherveniakova, T., & Christova, I. (2024). Resurgence of respiratory syncytial virus with dominance of RSV-B during the 2022–2023 season. Frontiers in Microbiology, 15. https://www.frontiersin.org/journals/microbiology/articles/10.3389/fmicb.2024.1376389

35. L, H. L., Ron, D., Yuan, Y., Manuel, B. C., Miroslava, B., A, M. S., J, M. W., J, Z. H., Dennis, B., Amy, G., Ulrika, W. H., S, M. V., Pin, R., Therese, T., E, A. M., Amanda, L., Pamela, G. M., & Tonya, V. (2022). Nirsevimab for Prevention of RSV in Healthy Late-Preterm and Term Infants. New England Journal of Medicine, 386(9), 837–846. 10.1056/NEJMoa2110275

36. Li, H., & Durbin, R. (2010). Fast and accurate long-read alignment with Burrows–Wheeler transform. Bioinformatics, 26(5), 589–595. 10.1093/bioinformatics/btp698

37. Li, H., Handsaker, B., Wysoker, A., Fennell, T., Ruan, J., Homer, N., Marth, G., Abecasis, G., Durbin, R., & Subgroup, 1000 Genome Project Data Processing. (2009). The Sequence Alignment/Map format and SAMtools. Bioinformatics, *25*(16), 2078–2079. 10.1093/bioinformatics/btp352

38. Li, Y., Johnson, E. K., Shi, T., Campbell, H., Chaves, S. S., Commaille-Chapus, C., Dighero, I., James, S. L., Mahé, C., Ooi, Y., Paget, J., van Pomeren, T., Viboud, C., & Nair, H. (2021). National burden estimates of hospitalisations for acute lower respiratory infections due to respiratory syncytial virus in young children in 2019 among 58 countries: a modelling study. The Lancet. Respiratory Medicine, 9(2), 175–185. 10.1016/S2213-2600(20)30322-2

39. Liao, Y., Smyth, G. K., & Shi, W. (2014). featureCounts: an efficient general purpose program for assigning sequence reads to genomic features. Bioinformatics, 30(7), 923–930. 10.1093/bioinformatics/btt656

40. Lydia, T., J, C. F. E., Lieselot, H., C, V. M., M, van B. G., J, W. E. J. H., P, M. D., & Philippe, L. (2013). The Comparative Genomics of Human Respiratory Syncytial Virus Subgroups A and B: Genetic Variability and Molecular Evolutionary Dynamics. Journal of Virology, 87(14), 8213–8226. 10.1128/jvi.03278-12

41. Majidian, S., Chalco, A., Zheng, X., Webby, R. J., Bowman, A. S., Poulson, R. L., Nemeth, N. M., Sedlazeck, F. J., & Agustinho, D. P. (2026). Rapid phylogenomic analysis for viral surveillance and metagenomic profiling with Omni2Tree. BioRxiv, 2026.04.29.721707. 10.64898/2026.04.29.721707

42. McFarland, E. J., Karron, R. A., Muresan, P., Cunningham, C. K., Libous, J., Perlowski, C., Thumar, B., Gnanashanmugam, D., Moye, J., Schappell, E., Barr, E., Rexroad, V., Fearn, L., Spector, S. A., Aziz, M., Cielo, M., Beneri, C., Wiznia, A., Luongo, C., … Buchholz, U. J. (2019). Live Respiratory Syncytial Virus Attenuated by M2-2 Deletion and Stabilized Temperature Sensitivity Mutation 1030s Is a Promising Vaccine Candidate in Children. The Journal of Infectious Diseases, 221(4), 534. 10.1093/INFDIS/JIZ603

43. McMorrow, M. L., Moline, H. L., Toepfer, A. P., Halasa, N. B., Schuster, J. E., Staat, M. A., Williams, J. V, Klein, E. J., Weinberg, G. A., Clopper, B. R., Boom, J. A., Stewart, L. S., Selvarangan, R., Schlaudecker, E. P., Michaels, M. G., Englund, J. A., Albertin, C. S., Mahon, B. E., Hall, A. J., … Curns, A. T. (2024). Respiratory Syncytial Virus-Associated Hospitalizations in Children <5 Years: 2016–2022. Pediatrics, 154(1), e2023065623. 10.1542/peds.2023-065623

44. NVSN. (2024, June). Pediatric Acute Respiratory Illness (ARI) Interactive Dashboard. https://www.cdc.gov/nvsn/php/ari-dashboard/

45. Peret, T. C., Hall, C. B., Schnabel, K. C., Golub, J. A., & Anderson, L. J. (1998). Circulation patterns of genetically distinct group A and B strains of human respiratory syncytial virus in a community. The Journal of General Virology, 79 ( *Pt 9*), 2221–2229. http://www.ncbi.nlm.nih.gov/pubmed/9747732

46. Perez, A., Lively, J. Y., Curns, A., Weinberg, G. A., Halasa, N. B., Staat, M. A., Szilagyi, P. G., Stewart, L. S., McNeal, M. M., Clopper, B., Zhou, Y., Whitaker, B. L., LeMasters, E., Harker, E., Englund, J. A., Klein, E. J., Selvarangan, R., Harrison, C. J., Boom, J. A., … McMorrow, M. (2022). Respiratory Virus Surveillance Among Children with Acute Respiratory Illnesses - New Vaccine Surveillance Network, United States, 2016-2021. MMWR. Morbidity and Mortality Weekly Report, *71*(40), 1253–1259. 10.15585/mmwr.mm7140a1

47. Pertea, G., & Pertea, M. (2020). GFF Utilities: GffRead and GffCompare [version 2; peer review: 3 approved]. F1000Research, *9*(304). 10.12688/f1000research.23297.2

48. Pierangeli, A., Midulla, F., Piralla, A., Ferrari, G., Nenna, R., Pitrolo, A. M. G., Licari, A., Marseglia, G. L., Abruzzese, D., Pellegrinelli, L., Galli, C., Binda, S., Cereda, D., Fracella, M., Oliveto, G., Campagna, R., Petrarca, L., Pariani, E., Antonelli, G., & Baldanti, F. (2024). Sequence analysis of respiratory syncytial virus cases reveals a novel subgroup -B strain circulating in north-central Italy after pandemic restrictions. Journal of Clinical Virology, 173, 105681. 10.1016/j.jcv.2024.105681

49. Piñana, M., González-Sánchez, A., Andrés, C., Vila, J., Creus-Costa, A., Prats-Méndez, I., Arnedo-Muñoz, M., Saubi, N., Esperalba, J., Rando, A., Nadal-Baron, P., Quer, J., González-López, J. J., Soler-Palacín, P., Martínez-Urtaza, J., Larrosa, N., Pumarola, T., & Antón, A. (2024). Genomic evolution of human respiratory syncytial virus during a decade (2013–2023): bridging the path to monoclonal antibody surveillance. Journal of Infection, 88(5), 106153. 10.1016/j.jinf.2024.106153

50. Rambaut, A., Drummond, A. J., Xie, D., Baele, G., & Suchard, M. A. (2018). Posterior Summarization in Bayesian Phylogenetics Using Tracer 1.7. Systematic Biology, 67(5), 901–904. 10.1093/SYSBIO/SYY032

51. Rios-Guzman, E., Simons, L. M., Dean, T. J., Agnes, F., Pawlowski, A., Alisoltanidehkordi, A., Nam, H. H., Ison, M. G., Ozer, E. A., Lorenzo-Redondo, R., & Hultquist, J. F. (2024). Deviations in RSV epidemiological patterns and population structures in the United States following the COVID-19 pandemic. Nature Communications, 15(1), 3374. 10.1038/s41467-024-47757-9

52. Schobel, S. A., Stucker, K. M., Moore, M. L., Anderson, L. J., Larkin, E. K., Shankar, J., Bera, J., Puri, V., Shilts, M. H., Rosas-Salazar, C., Halpin, R. A., Fedorova, N., Shrivastava, S., Stockwell, T. B., Peebles, R. S., Hartert, T. V., & Das, S. R. (2016). Respiratory Syncytial Virus whole-genome sequencing identifies convergent evolution of sequence duplication in the C-terminus of the G gene. Scientific Reports, 6. 10.1038/srep26311

53. Shepard, S. S., Meno, S., Bahl, J., Wilson, M. M., Barnes, J., & Neuhaus, E. (2016). Viral deep sequencing needs an adaptive approach: IRMA, the iterative refinement meta-assembler. BMC Genomics, 17(1), 708. 10.1186/s12864-016-3030-6

54. Simões, E. A. F., Forleo-Neto, E., Geba, G. P., Kamal, M., Yang, F., Cicirello, H., Houghton, M. R., Rideman, R., Zhao, Q., Benvin, S. L., Hawes, A., Fuller, E. D., Wloga, E., Pizarro, J. M. N., Munoz, F. M., Rush, S. A., McLellan, J. S., Lipsich, L., Stahl, N., … Sivapalasingam, S. (2021). Suptavumab for the Prevention of Medically Attended Respiratory Syncytial Virus Infection in Preterm Infants. Clinical Infectious Diseases, 73(11), e4400–e4408. 10.1093/cid/ciaa951

55. Sondlane, H., Ogunbayo, A., Donato, C., Mogotsi, M., Esona, M., Hallbauer, U., Bester, P., Goedhals, D., & Nyaga, M. (2024). Whole genome molecular analysis of respiratory syncytial virus pre and during the COVID-19 pandemic in Free State province, South Africa. Virus Research, 347, 199421. 10.1016/j.virusres.2024.199421

56. Stamatakis, A. (2014). RAxML version 8: a tool for phylogenetic analysis and post-analysis of large phylogenies. Bioinformatics, 30(9), 1312–1313. 10.1093/bioinformatics/btu033

57. Tan, L., Lemey, P., Houspie, L., C. Viveen, M., Jansen, N. J. G., van Loon, A. M., Wiertz, E., van Bleek, G. M., Martin, D. P., & Coenjaerts, F. E. (2012). Genetic Variability among Complete Human Respiratory Syncytial Virus Subgroup A Genomes: Bridging Molecular Evolutionary Dynamics and Epidemiology. PLOS ONE, 7(12), e51439-. 10.1371/journal.pone.0051439

58. Terstappen, J., Hak, S. F., Bhan, A., Bogaert, D., Bont, L. J., Buchholz, U. J., Clark, A. D., Cohen, C., Dagan, R., Feikin, D. R., Graham, B. S., Gupta, A., Haldar, P., Jalang’o, R., Karron, R. A., Kragten, L., Li, Y., Löwensteyn, Y. N., Munywoki, P. K., … Mazur, N. I. (2024). The respiratory syncytial virus vaccine and monoclonal antibody landscape: the road to global access. The Lancet Infectious Diseases. 10.1016/S1473-3099(24)00455-9

59. Trento, A., Viegas, M., Galiano, M., Videla, C., Carballal, G., Mistchenko, A. S., & Melero, J. A. (2006). Natural history of human respiratory syncytial virus inferred from phylogenetic analysis of the attachment (G) glycoprotein with a 60-nucleotide duplication. Journal of Virology, 80(2), 975–984. 10.1128/JVI.80.2.975-984.2006

60. Turakhia, Y., Thornlow, B., Hinrichs, A. S., De Maio, N., Gozashti, L., Lanfear, R., Haussler, D., & Corbett-Detig, R. (2021). Ultrafast Sample placement on Existing tRees (UShER) enables real-time phylogenetics for the SARS-CoV-2 pandemic. Nature Genetics, 53(6), 809–816. 10.1038/s41588-021-00862-7

61. Ujiie, M., Tsuzuki, S., Nakamoto, T., & Iwamoto, N. (2021). Resurgence of Respiratory Syncytial Virus Infections during COVID-19 Pandemic, Tokyo, Japan. Emerging Infectious Disease Journal, 27(11), 2969. 10.3201/eid2711.211565

62. Venter, M., Madhi, S. A., Tiemessen, C. T., & Schoub, B. D. (2001). Genetic diversity and molecular epidemiology of respiratory syncytial virus over four consecutive seasons in South Africa: identification of new subgroup A and B genotypes. The Journal of General Virology, 82(Pt 9), 2117–2124. http://www.ncbi.nlm.nih.gov/pubmed/11514720

63. Wang, S., Sundaram, J. P., & Stockwell, T. B. (2012). VIGOR extended to annotate genomes for additional 12 different viruses. Nucleic Acids Research, 40(W1), W186–W192. 10.1093/nar/gks528

64. Wood, D. E., Lu, J., & Langmead, B. (2019). Improved metagenomic analysis with Kraken 2. Genome Biology, 20(1), 257. 10.1186/s13059-019-1891-0

65. Yu, G., Smith, D. K., Zhu, H., Guan, Y., & Lam, T. T. Y. (2017). ggtree: an r package for visualization and annotation of phylogenetic trees with their covariates and other associated data. Methods in Ecology and Evolution, 8(1), 28–36. 10.1111/2041-210X.12628;WEBSITE:WEBSITE:BESJOURNALS;WGROUP:STRING:PUBLICATION

66. Yu, J.-M., Fu, Y.-H., Peng, X.-L., Zheng, Y.-P., & He, J.-S. (2021). Genetic diversity and molecular evolution of human respiratory syncytial virus A and B. Scientific Reports, 11(1), 12941. 10.1038/s41598-021-92435-1

67. Zhu, Q., McAuliffe, J. M., Patel, N. K., Palmer-Hill, F. J., Yang, C., Liang, B., Su, L., Zhu, W., Wachter, L., Wilson, S., MacGill, R. S., Krishnan, S., McCarthy, M. P., Losonsky, G. A., & Suzich, J. A. (2011). Analysis of respiratory syncytial virus preclinical and clinical variants resistant to neutralization by monoclonal antibodies palivizumab and/or motavizumab. The Journal of Infectious Diseases, 203(5), 674–682. 10.1093/infdis/jiq100

