## Supplementary figures and images for "Evolutionary dynamics of Respiratory Syncytial Virus in pre-pandemic, pandemic, and post-pandemic periods in Houston, Texas, USA"

### Supplemental Figure 1a

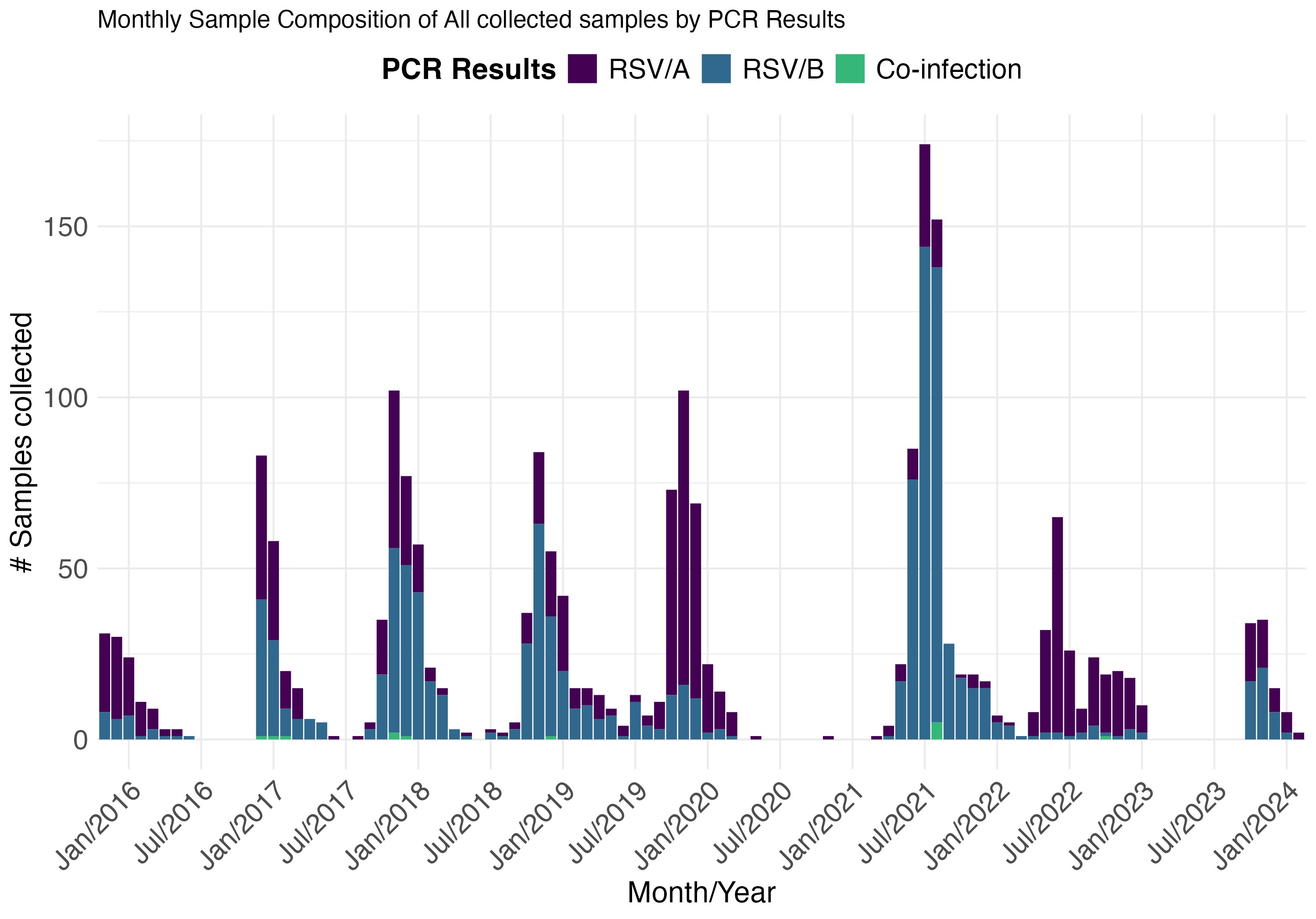

### Supplemental Figure 1b

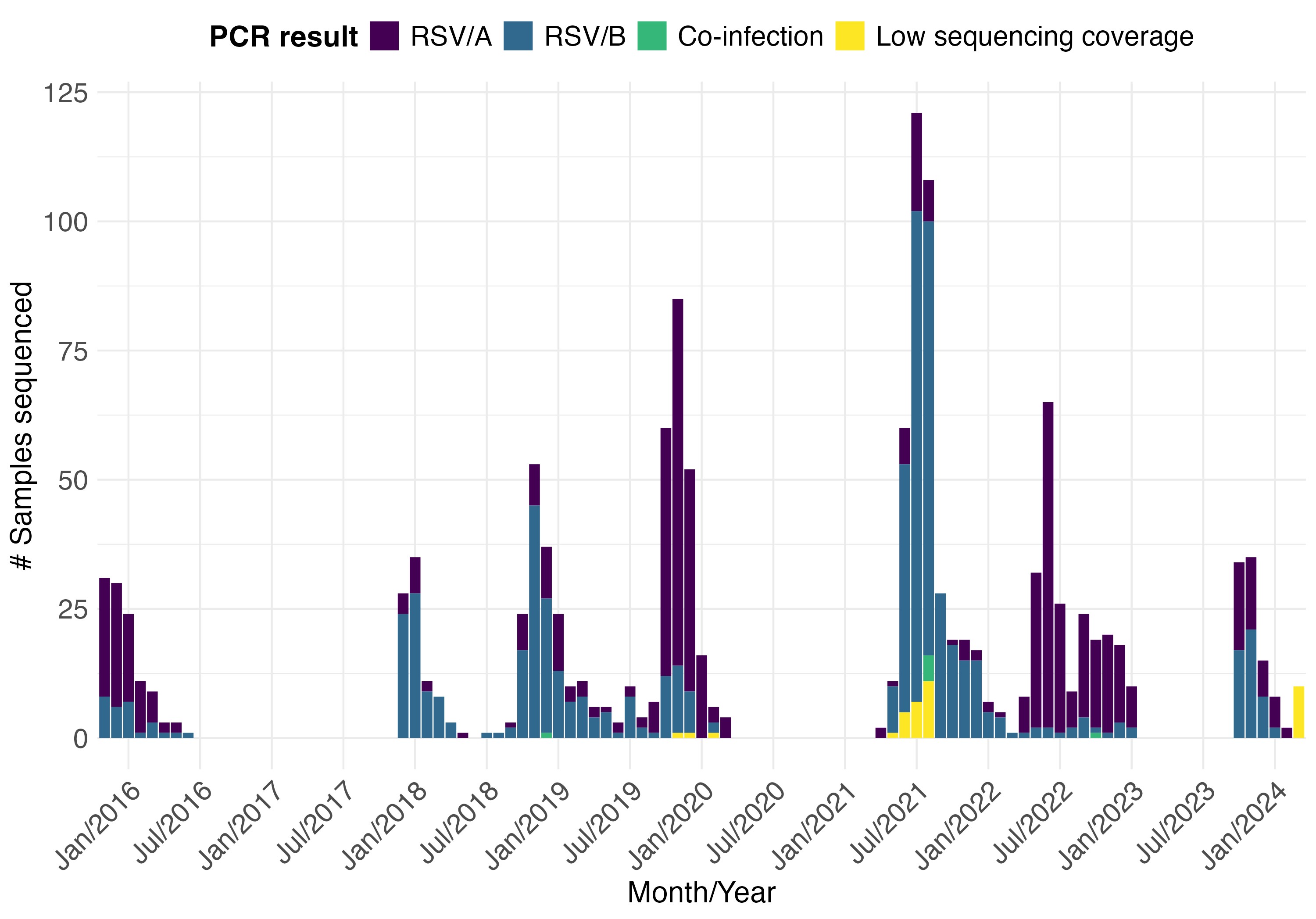

### Supplemental Figure 2

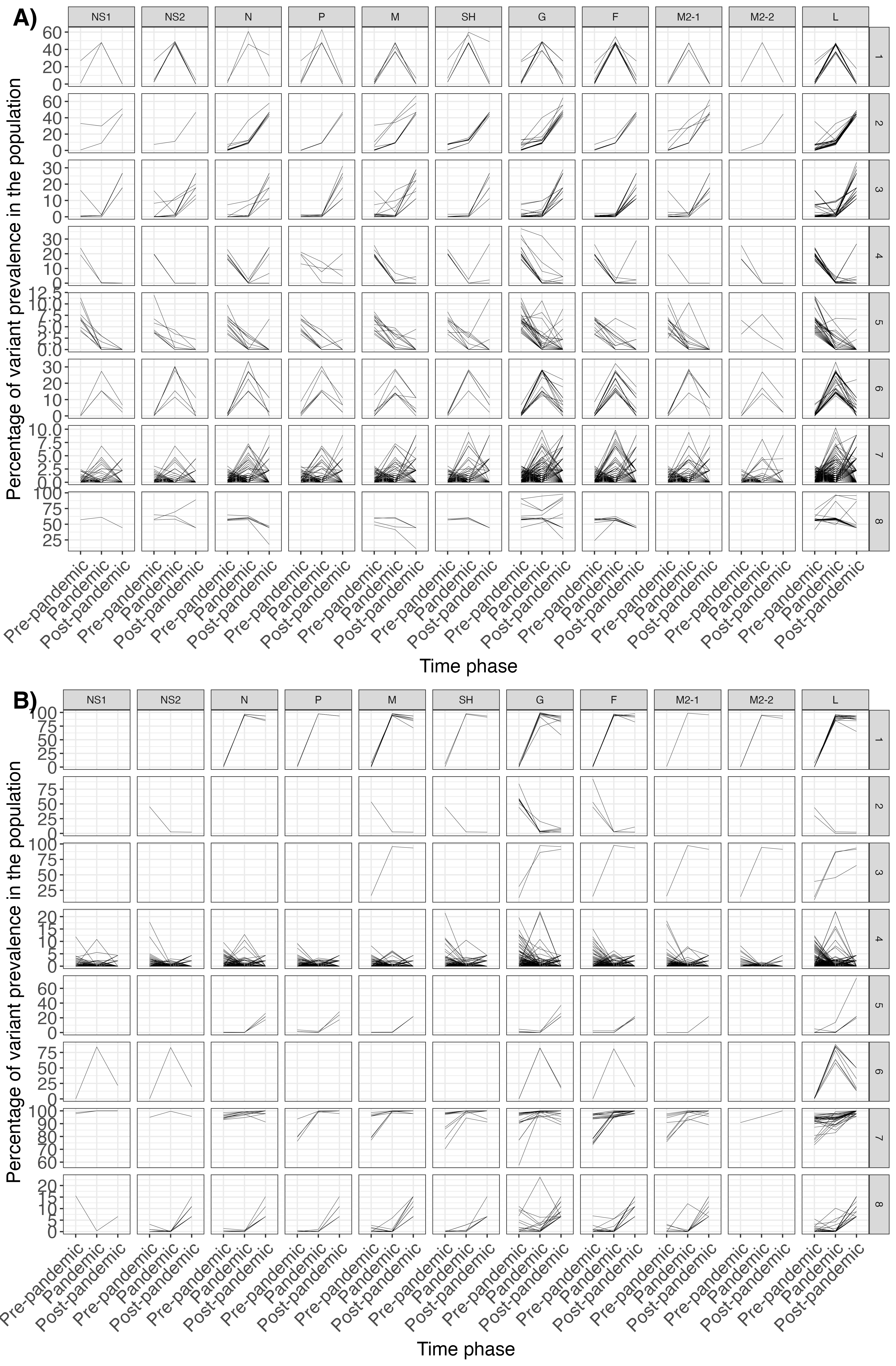

### Supplemental Figure 3

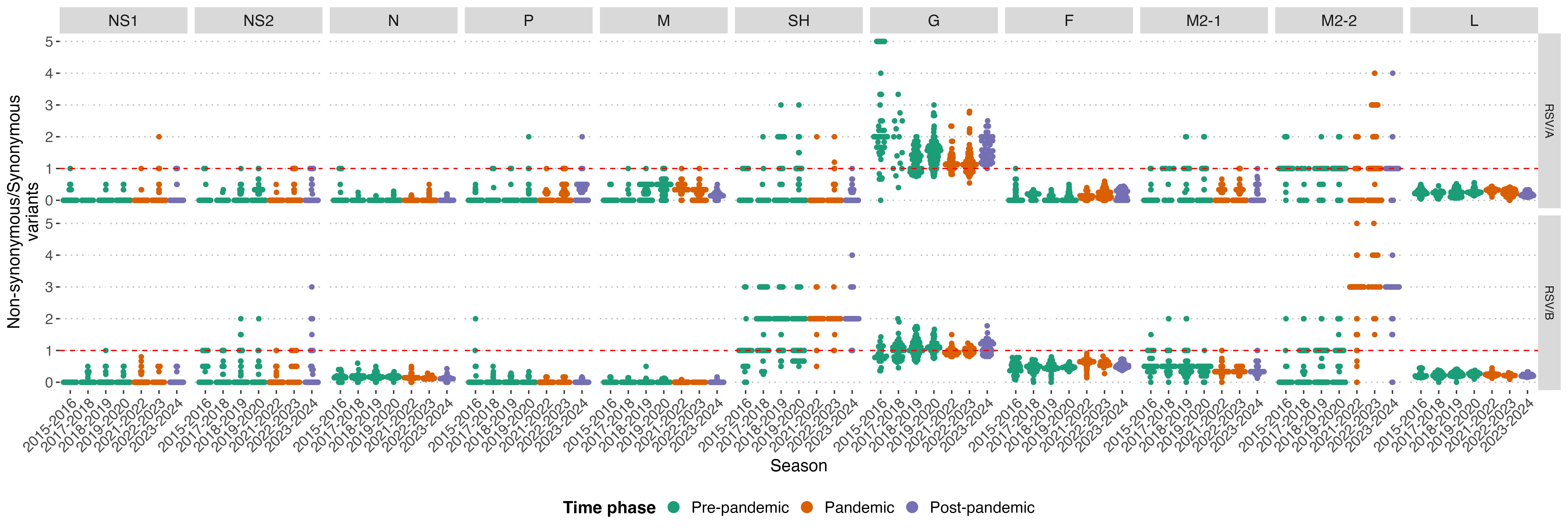

### Supplemental Figure 4

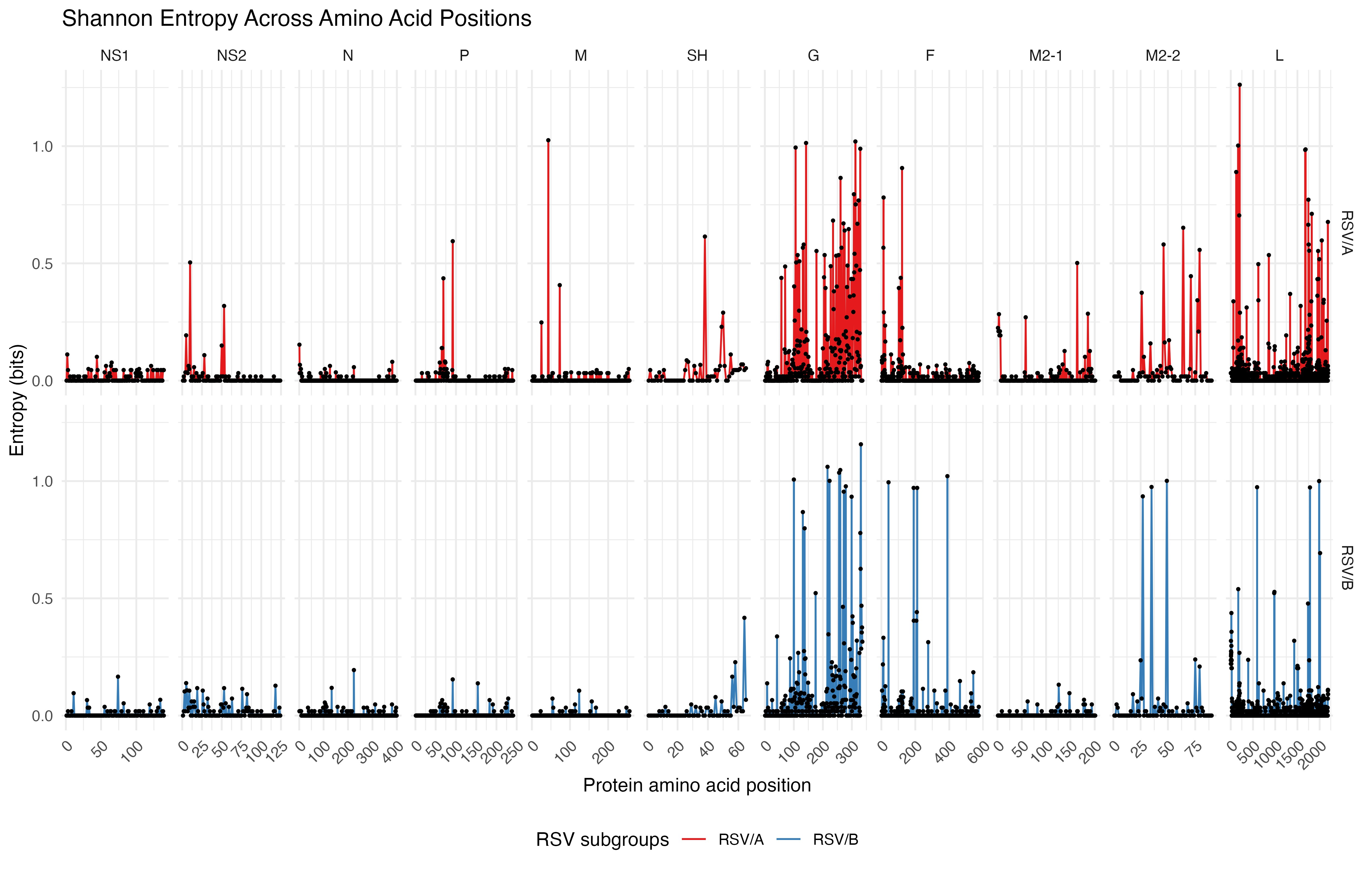
